# Functional Ambidexterity of an Ancient Nucleic Acid-Binding Domain

**DOI:** 10.1101/2023.03.06.531422

**Authors:** Orit Weil-Ktorza, Dragana Despotović, Yael Fridmann-Sirkis, Segev Naveh-Tassa, Yaacov Levy, Norman Metanis, Liam M. Longo

## Abstract

Homochirality of biopolymers emerged early in the history of life on Earth, nearly 4 billion years ago. Whether the establishment of homochirality was the result of abiotic physical and chemical processes, or biological selection, remains unknown. However, given that significant events in protein evolution predate the last universal common ancestor, the history of homochirality may have been written into some of the oldest protein folds. To test this hypothesis, the evolutionary trajectory of the ancient and ubiquitous helix-hairpin-helix (HhH) protein family was analyzed for functional robustness to total chiral inversion of just one binding partner. Against expectations, functional ‘ambidexterity’ was observed across the entire trajectory, from phase separation of HhH peptides with RNA to dsDNA binding of the duplicated (HhH)_2_-Fold. Moreover, dissociation kinetics, mutational analysis, and molecular dynamics simulations revealed significant overlap between the binding modes of a natural and a mirror-image protein to natural dsDNA. These data suggest that the veil between worlds with alternative chiral preferences may not be as impenetrable as is often assumed, and that the HhH protein family is an intriguing exception to the dogma of reciprocal chiral substrate specificity proposed by Milton and Kent (Milton *et al*. Science 1992).

## Introduction

Life on earth is characterized by exquisite homochirality: With very few exceptions, proteins and peptides are composed of *L*-amino acids (except for glycine, which is achiral) while nucleic acids (RNA and DNA) are derived from *D*-ribose. The history of homochirality is murky, no doubt because its origins predate the last universal common ancestor (LUCA), which likely arose more than 3.5 billion years ago.^1, 2, 3^ Consequently, theories regarding the emergence and evolution of homochirality remain speculative and tend to emphasize chemical and physical processes over biological selection.^4^ The benefits of chiral control over amino acid incorporation, however, are clear: By stabilizing and regularizing the conformations of proteins, especially protein secondary structures,^5^ homochirality laid the groundwork for the emergence of folded protein domains.

Current evidence suggests that the LUCA already had an assortment of folded protein domains with complex structure^6^, indicating that significant events in protein evolution predate the LUCA itself. The helix-hairpin-helix (HhH) motif^7^, for example, is an ancient nucleic acid binding element that is ubiquitous across the tree of life and present in ribosomal protein S13.^8^ It binds the phosphate backbone of RNA and DNA via the N-terminus of an α-helix, a primitive binding mode^9^, and was identified as among the primordial peptides around which modern domains condensed.^10^ Moreover, the ancestral sequence of the HhH binding loop, PGIGP, is a palindrome composed of ancient amino acids.^11, 12^ Duplication of the HhH motif yields a globular domain called the (HhH)_2_-Fold that binds to the minor groove of dsDNA.^8^ We have previously used the HhH motif to study the early functional evolution of nucleic acid binding, where we observed a transition from a simple phase separating peptide to a structured domain with specific dsDNA binding functionality (**Figure 1**).^13^ The (HhH)_2_-Fold, like other putative ancient folds^14, 15^, can be encoded by restricted alphabets biased for ancient amino acids^16^, including ornithine as a precursor to arginine.^13^ More recently, we have demonstrated that dimerization of the HhH motif, likely to form an (HhH)_2_-Fold, is promoted upon RNA binding and occurs within peptide-RNA coacervates.^17^ Given the deep antiquity of the HhH motif and the derived (HhH)_2_-Fold,^7, 8, 10^ is it possible that these structures were shaped by the early evolution of homochirality? How could indications of such a history be uncovered?

**Figure 1.**
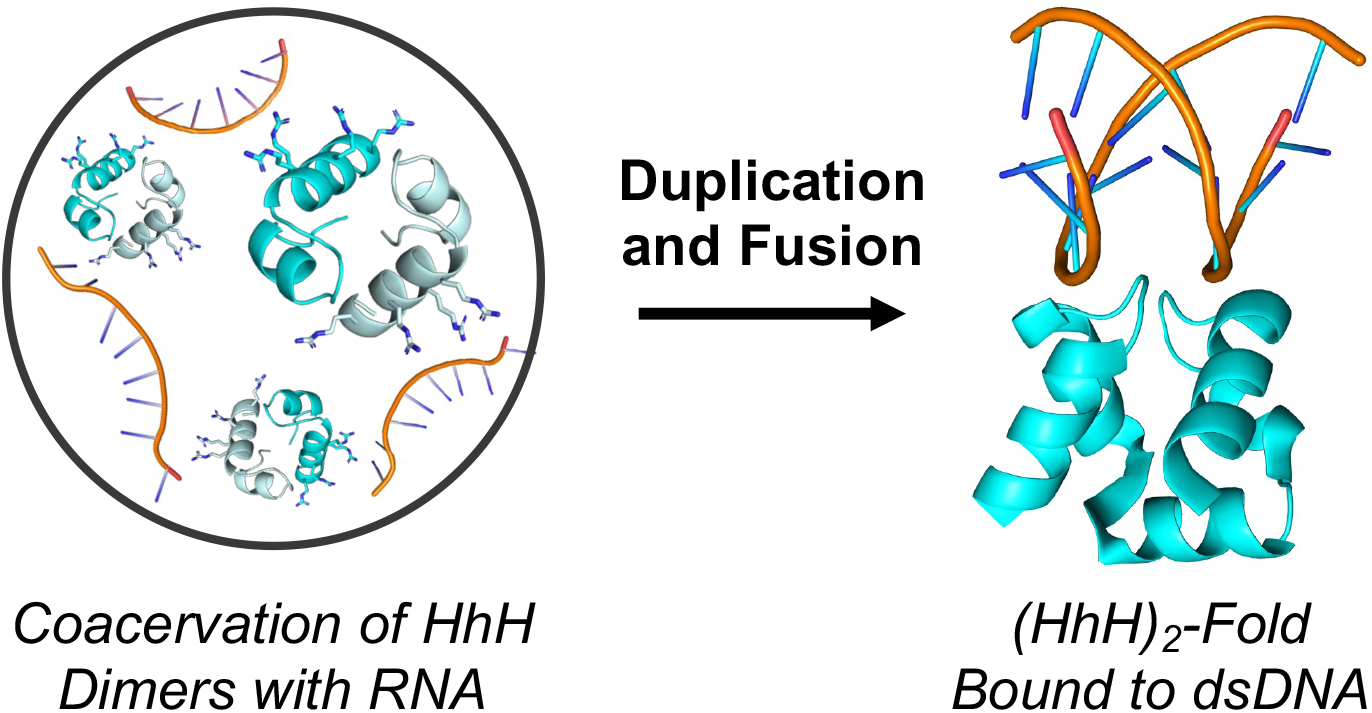
Transition from a dimerizing HhH motif that forms coacervates with RNA to a monomeric (HhH)_2_-Fold that binds to the minor groove of dsDNA binding (PDB ID 1c7y). Structure figures made using PyMol.

First, it is necessary to consider the consequences of chiral inversion in modern biology, which is typically associated with a catastrophic loss of functionality, as famously demonstrated by Milton and Kent.^18^ Indeed, this outcome is so reliable that several technologies have been developed to take advantage of it, such as Spieglemers^19^ (aptamers with inverted chirality) and mirror-image phage display.^20, 21, 22, 23^ Therapeutic compounds with inverted chirality are exceptionally resistant to endogenous nucleases and proteases, resulting in extended *in vivo* half-lifes.^24^ As such, mirror image enzymes have also been employed to degrade environmental pollution by achiral plastic substrates because these enzymes have superior biostability.^25^ Even coacervate formation has been demonstrated to be sensitive to chiral inversion of just one of the interacting partners.^26^ One might reasonably predict, then, that total chiral inversion of the HhH motif or the (HhH)_2_-Fold will disrupt their respective functionalities of phase separation with RNA and binding to dsDNA. After all, both proteins and nucleic acids are composed of chiral residues (*i.e*., *L*-amino acids and *D*-ribose, respectively) that fold into chiral conformers (*e.g*., right-handed □-helix and right-handed double helix) (**Figure 1**).

Following the previously reported trajectory of ancestor reconstruction and experimental deconstruction,^13^ we now report that the function of both the HhH motif and the (HhH)_2_-Fold are surprisingly robust to total chiral inversion. In the case of the (HhH)_2_-Fold binding to dsDNA, entropy mutations in the canonical binding loop (*i.e*., PGIGP → GGGGG) were more disruptive to dsDNA binding than was total chiral inversion of the protein. Molecular dynamics simulations revealed that, remarkably, the residues that mediate binding of *L*-protein to *D*-dsDNA are largely retained in the *D*-protein:*D*-dsDNA (mirror protein:natural dsDNA) binding mode. We must now grapple with the question of whether functional ‘ambidexterity’ of an ancient nucleic-acid binding domain may report on the early history of homochirality.

## Materials and Methods

### Total Protein Synthesis

Proteins and peptides characterized in this study were prepared by total chemical synthesis using solid-phase peptide synthesis (SPPS)^27^ and native chemical ligation (NCL)^28^ followed by desulfurization of the Cys residue at the ligation site, as described previously^13, 17^ (see **SI** for a detailed protocol and **Figures S1-S19** and **Schemes S1 and S2**). Construct names and sequences are given in **Table 1**.

**Table 1.**
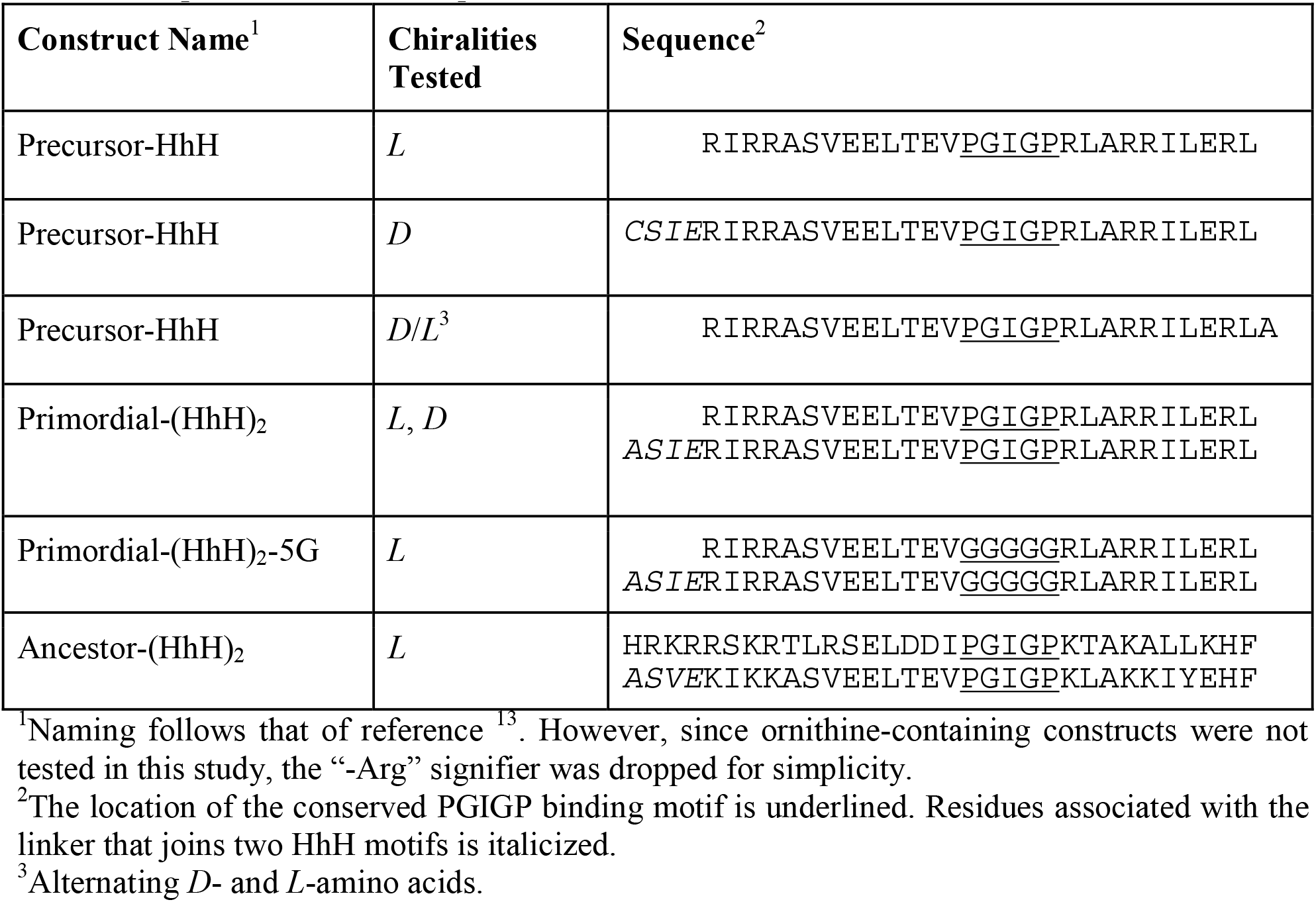
Peptide and Protein Sequences.

### Circular Dichroism

Circular dichroism (CD) spectra were collected on a Chirascan CD spectrometer (Applied Photophysics) with a 1-mm pathlength quartz cuvette. Samples containing 10 μM *L*-Precursor-HhH, *D*-Precursor-HhH or *D/L*-Precursor-HhH were measured in 5 mM Tris·HCl, 50 mM NaCl, pH 7.5 with either 0% or 20% (v/v) trifluoroethanol (TFE). Samples containing 5 μM *L-* Primordial-(HhH)_2_, *D*-Primordial-(HhH)_2_ or *L*-Primordial-(HhH)_2_-5G were measured in 5 mM Tris·HCl, 500 mM NaCl, 10 mM MgCl_2_, 5 mM CaCl_2_, pH 7.5. All spectra were collected from 195 to 260 nm with a data pitch of 1 nm at room temperature, 3 sec signal averaging per data point, and a slit width of 1 nm. The photomultiplier tube voltage during measurement was kept below 700 V, and data points exceeding this value were discarded. All reported spectra have had the spectrum of the buffer subtracted.

### Phase Separation

Peptides and polyuridylic acid (polyU; Sigma-Aldrich, P9528) were dissolved in Milli-Q water (Millipore Sigma). Peptide concentrations were measured using the Pierce BCA Protein Assay Kit (Thermo Fisher Scientific). Stock solutions of 10 mg/mL polyU and 500 mM MES pH 5.6 were prepared. Phase separation was induced by mixing the peptide and polyU solutions. The final composition of the phase separation reaction mixture was 50 mM MES pH 5.6, 1.0 mg/mL polyU, and 300 μM peptide. Typically, 3 μL of the phase separation reaction mixture was loaded onto slides (24 × 40 mm, 0.13-0.16 mm thick) and observed using an Eclipse TI-E Nikon inverted microscope (Nikon Instruments Inc., Melville, NY) with an oil-immersion 100× objective (Plan Apo, 100×/1.40 oil). Images were acquired with a cooled electron-multiplying charge-coupled device camera (IXON ULTRA 888, Andor). Pictures were analyzed using the Fiji platform.^29^

### NanoSight

Coacervates were prepared as described in the previous section. The final composition of the phase separation reaction mixture was 50 mM MES pH 5.6, 1.0 mg/mL polyU, and 300 μM peptide. Upon phase separation, samples were diluted 500-fold in 50 mM MES pH 5.6 and the size distribution and concentration of the droplets were measured by a NanoSight NS300 (Malvern Panalytical Ltd, UK) using a 405 nm laser and an sCMOS camera. The camera level was increased until all particles were distinctly visible (level 11). For each measurement, five 1-min videos were captured at a cell temperature of 25 °C. The videos were analyzed by the built-in NanoSight Software (NTA 3.4 Build 3.4.003) with a detection threshold of 2.

### Surface Plasmon Resonance

Binding of biotinylated 29-bp *D*-dsDNA and *L*-dsDNA (the mirror-image DNA derived from *L*-ribose) as well as 29-bp *D*-ssDNA (see **Table S1** for DNA sequences) was monitored by surface plasmon resonance (SPR) on a Biacore S200 system (Cytiva, Sweden). Since the (HhH)_2_-Fold variants are positively charged at neutral pH, a C1 chip (S-Series Cytiva, Sweden), which carries less negative charge than the standard CM5 chip, was used. Streptavidin was conjugated to the chip surface using EDC/NHS chemistry, as outlined in the C1 sensor chip manual.

Approximately 2000 RU (Chip 1, **Main Text**; buffer pH 3.8) or 700 RU (Chip 2, **SI**; buffer pH 4.6) of streptavidin was covalently conjugated to the chip surface and then blocked by injecting 1 M ethanolamine pH 8.0 for 5 min. Subsequently, 405 RU of *D*-dsDNA, 423 RU of *L*-dsDNA and 210 RU of *D*-ssDNA (Chip 1) or 95 RU of *D*-dsDNA, 97 RU of *L*-dsDNA and 65 RU of *D*-ssDNA (Chip 2) were stably associated to the surface of one channel. Before data collection, a normalization cycle followed by three priming cycles were run to stabilize the instrument. Binding assays were performed in SPR binding buffer (50 mM Tris, 150 mM NaCl, 0.05% Tween-20, pH 7.5) with a flow rate of 20 μL/min at 25 °C. Regeneration of the chip surface was achieved by a 60 s injection of 2 M NaCl in water. Reported sensorgrams were double subtracted: First, by the background binding of the analyte to a streptavidin-conjugated control channel and then by the average of 2 buffer injection runs.

### Microscale Thermophoresis (MST)

Protein-DNA interactions were analyzed by microscale thermophoresis. Experiments were performed with 25 nM of synthetic, Cy5-labelled (HhH)_2_-Fold proteins, which were prepared by coupling Cy5-NHS ester to the N-terminus of synthetic proteins (note that there are no Lys residues present in these sequences). Experiments were carried out in a microMonolith NT.115 Blue/Red (NanoTemper Technologies) at 25 °C. Labelled peptides were mixed with serially diluted DNA samples in 50 mM Tris, 150 mM NaCl, 0.05% Tween-20, pH 7.5 in premium capillaries (NanoTemper Technologies) at 40% MST power. Dissociation constants (*K_D_*) could not be calculated as binding is non-specific for the minor groove of dsDNA and, as a result, one strand of dsDNA has many degenerate, overlapping binding sites. See **Table S1** for DNA sequences.

### Molecular Dynamics (MD) Simulations

An atomistic model of *L*-Primordial-(HhH)_2_ was made using the ColabFold implementation of AlphaFold2.^30^ This updated model was in good agreement with an earlier homology model from SWISS-MODEL,^17, 31^ particularly in the region of the nucleic acid binding loops. The initial dsDNA binding orientation for the *L*-Primordial-(HhH)_2_ was estimated by alignment to the natural (HhH)_2_-Fold:dsDNA complex in PDB ID 1c7y using the *cealign* algorithm in PyMOL (pymol.org). The DNA sequences were taken from chains C and G of PDB ID 1c7y and elongated to 21 bp by repetition (see **Table S1** for DNA sequences). *D*-Primordial-(HhH)_2_ was generated using DStabilize,^32^ and *L*-Primordial-(HhH)_2_-5G (**Table 1**) was generated using PyMOL mutagenesis of *L*-Primordial-(HhH)_2_. MD simulations were performed using the CHARMM36 force field^33^ in GROMACS 2022.1.^34^ Each system was solvated in a water box (TIP3P) with 0.125 M NaCl, where the ions also served to render each system neutral. Each system was minimized with the steepest descent method, followed by equilibration with the NVT and NPT regimes.^35^ Position restraints were applied to both ends of each dsDNA strand (one base pair) during production runs, which lasted 1 μs at a temperature of 300 K and were repeated three times for each system.

## Results

### Coacervation of an HhH Motif with RNA is Functionally Ambidextrous

Although coacervate formation with RNA can be achieved by simple peptides,^36^ we have previously demonstrated that the amino acid composition of the *L*-Precursor-HhH peptide (**Table 1**) is not sufficient for droplet formation^13^: shuffling the sequence of *L*-Precursor-HhH -- either completely or preserving the positions of the basic amino acids -- resulted in a polypeptide that forms insoluble aggregates when mixed with *D*-polyU.^13^ Subsequent studies demonstrated that binding to RNA and coacervation were associated with dimerization of the peptide.^17^ To explore the functional profile of Precursor-HhH and its degree of ambidexterity in greater detail, *L*- and *D*-Precursor-HhH were synthesized (**Table 1**; see SI for details). Note that all chiral centers, including those within amino acid side chains (*i.e*., Thr and Ile), were inverted (mirror-image amino acids). The resulting CD spectra (**Figure 2A**) are indicative of unfolded, mirror image peptides. The addition of 20% trifluoroethanol, which induces α-helix formation,^37^ yielded mirror image spectra with characteristic peaks at 208 and 222 nm (**Figure 2A**).

**Figure 2.**
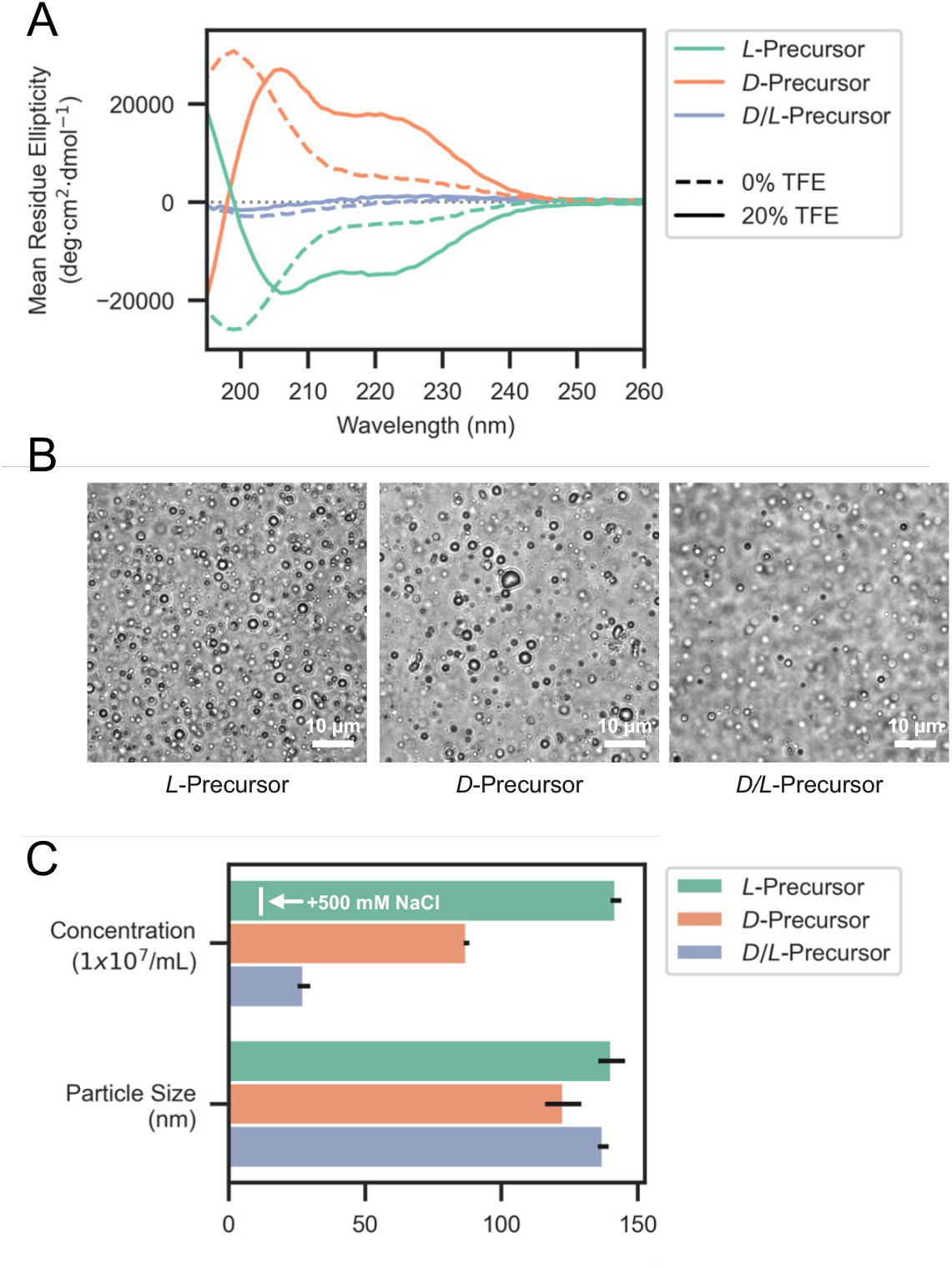
Coacervation of Precursor-HhH with PolyU in response to chiral inversion. **A**. Circular dichroism spectra of *D*- and *L*-Precursor-HhH indicates that the peptides are largely unfolded and are mirror images of each other. Addition of trifluoroethanol (TFE) induces α-helices in both cases, as indicated by the development of peaks at 208 nm and 222 nm. The *D*/*L*-Precursor-HhH peptide, which is comprised of amino acids of alternating chirality, does not exhibit significant circular dichroism signal alone or in the presence of TFE. **B**. Light micrographs of coacervates formed by 300 μM peptide and 1 mg/mL PolyU. *D*/*L*-Precursor-HhH consistently exhibited reduced coacervate formation. **C**. Nanoparticle tracking analysis of droplets. To achieve appropriate droplet concentrations, samples of 300 μM peptide and 1 mg/mL PolyU were diluted by a factor of 500 in buffer. Despite the dramatic change in component concentrations, droplets were sufficiently stable for measurement. Addition of 500 mM NaCl to the sample dramatically reduced the particle count, confirming that the observed particles were in fact coacervates. Although the sizes of the droplets in each sample were approximately the same, *D*/*L*-Precursor-HhH exhibited reduced droplet counts, consistent with the micrograph images and with a previous analysis that identifies α-helix formation and dimerization as being coupled to RNA binding and coacervation^17^.

Upon addition of *D*-PolyU, both *L*-Precursor-HhH and *D*-Precursor-HhH formed coacervates (**Figure 2B** and **C**). For a more quantitative measurement of coacervation forming potential, particle tracking analyses of coacervate-containing solutions was performed. To achieve the correct number of droplets in frame at any given time, a 500-fold dilution of previously-formed coacervates was imaged. Despite these concentrations being significantly lower than the concentrations used to drive coacervation, the droplets themselves were observed to be stable post-dilution, though of generally smaller size. To confirm that the detected particles were in fact droplets, 500 mM NaCl, which near-completely disrupts coacervation of Precursor-HhH, was added to a solution of droplets to serve as a control (**Figure 2C**). As expected, the resulting droplet counts were dramatically reduced (12.6 ± 0.3 × 10^7^ versus 142 ± 2.0 × 10^7^), though still slightly higher than either peptide (5.0 ± 1.1 × 10^7^) or *D*-polyU (1.4 ± 0.7 × 10^7^) alone. Estimates of the number of droplets formed by each peptide in the presence of *D*-polyU suggest that both the *L*- and *D*-peptides are similar in their droplet-forming potential, indicating that coacervation is robust to chiral inversion of just one partner and thus functionally ambidextrous.

We have previously shown by electron paramagnetic resonance (EPR) analysis that dimerization, RNA binding, and coacervation are linked processes.^17^ If true, phase separation potential may therefore depend in part on dimerization and α-helical folding. The α-helical folding potential of Precursor-HhH was abolished by alternating the chirality of every other amino acid^5^. The resulting construct, *D/L*-Precursor-HhH (**Table 1**, and SI for details), exhibited almost no circular dichroism with or without 20% trifluoroethanol, as it is expected (**Figure 2A**). *D/L*-Precursor-HhH also formed coacervates upon addition of polyU (**Figure 2B** and **C**) but did so more weakly than either the *D*- or *L*-homochiral variants. Coacervation is therefore not strictly dependent on Precursor-HhH α-helical folding, though it would seem that the emergence of α-helical folding can improve coacervation potential, as previously hypothesized.^17^

### Chiral Inversion of Primordial-(HhH)_2_

While the interactions that mediate coacervation are likely to be relatively nonspecific, the binding of Primordial-(HhH)_2_ to the minor groove of dsDNA (**Figure 1**) is highly dependent on the conformation of the protein itself, and should therefore be sensitive to chiral inversion. After all, both biopolymers adopt higher order chiral conformations. To probe the effect of chiral inversion, *D*-Primordial-(HhH)_2_ and *L*-Primordial-(HhH)_2_ were prepared by total chemical protein synthesis (**Figure 3A**; see **SI** for details). Circular dichroism spectra (**Figure 3B**) indicate that both *D*- and *L*-proteins fold into □-helical structures, with peaks around 208 and 222 nm. Just like the proteins themselves, and as expected, the CD spectra are mirror images of each other: *L*-Primordial-(HhH)_2_ has negative peaks (consistent with *L*-amino acids and right-handed □-helices) and *D*-Primordial-(HhH)_2_ has positive peaks (consistent with *D*-amino acids and left-handed □-helices). Minor differences between the spectra are likely due to the reduced purity of *D*-amino acid reagents and errors in concentration determination^38, 39^.

**Figure 3.**
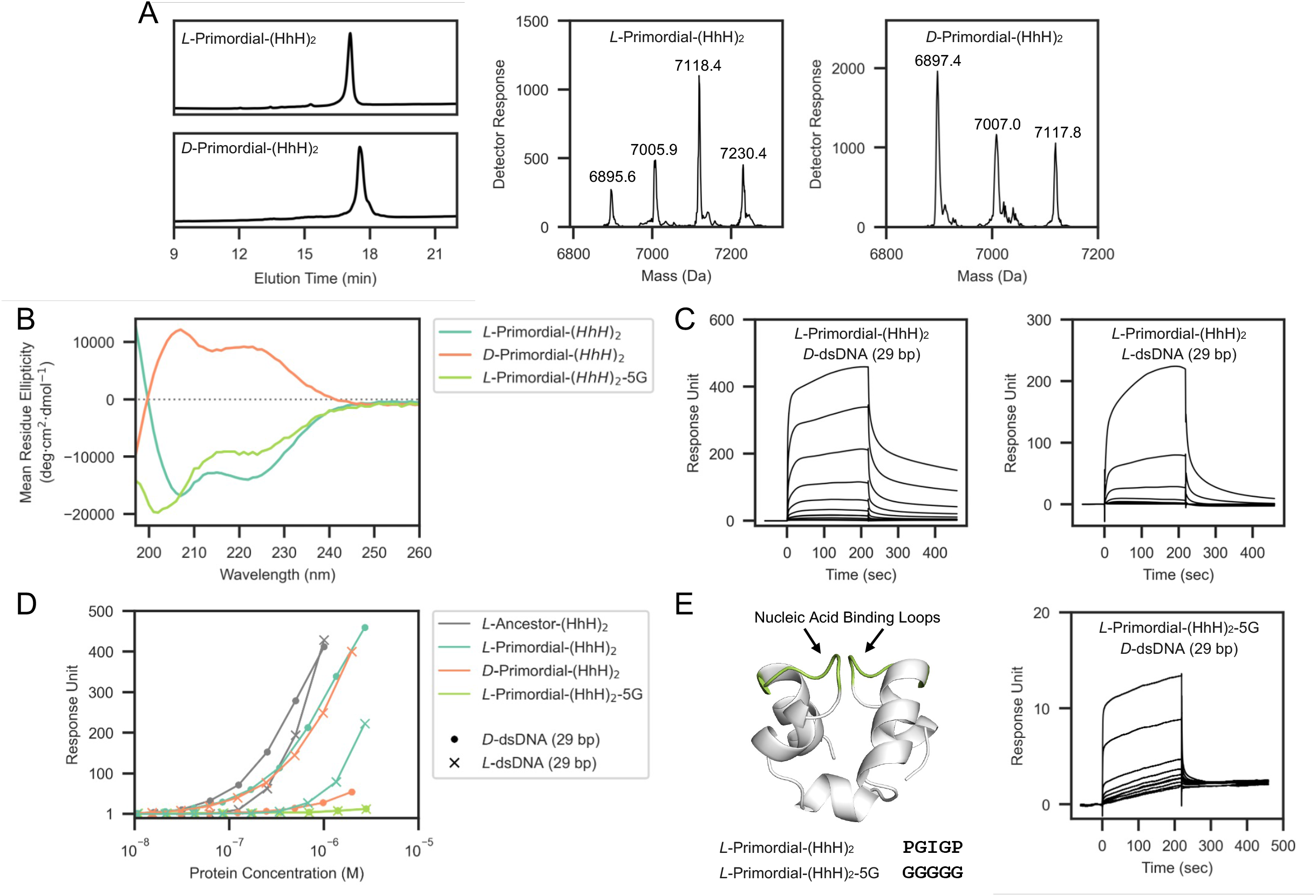
dsDNA binding of Primordial-(HhH)_2_ in response to chiral inversion. **A**. *L*- and *D*-Primordial-(HhH)_2_ was synthesized by a single native chemical ligation reaction between two peptide segments, followed by a desulfurization step to convert the Cys residue at the ligation site back to Ala (as described previously^13,17^, see **SI** for details). **B**. Circular dichroism spectra of *L*-Primordial-(HhH)_2_ and *D*-Primordial-(HhH)_2_ exhibit peaks around 208 nm and 222 nm, which are hallmarks of α-helical structure. The spectra are approximately symmetric about the x-axis, consistent with total chiral inversion and α-helices of opposite handedness. **C**. SPR sensorgrams demonstrating the binding of *L*-Primordial-(HhH)_2_ to immobilized *L*-dsDNA and *D*-dsDNA (protein concentration range 5.3 nM - 2.7 μM). **D**. Steady state analysis of SPR binding data. The ability of both Primordial-(HhH)_2_ and the more modern *L*-Ancestor-(HhH)_2_ to bind to either chiral form of dsDNA suggests that functional ambidexterity is a property of the fold itself, and not related to the highly simplified sequence of Primordial-(HhH)_2_. The amount of protein bound at state binding was approximated to be 4 s prior to the end of analyte injection, 216 s in total. An analysis of the dissociation rates is presented in **Figures S20** and **S21**. See **Figure S22** for replicate SPR binding data. **E**. The inactivation mutant *L*-Primordial-(HhH)_2_-5G, in which the canonical PGIGP binding motif (green) has been mutated to GGGGG (left). This mutant is less well-folded than the wild-type proteins (**Panel B**), and has greatly impaired binding affinity to dsDNA of either chirality (right, **Panel D**; concentration range 5.4 nM - 2.8 μM).

### dsDNA binding by Primordial-(HhH)2 is Functionally Ambidextrous

Binding of *D*-Primordial-(HhH)_2_ and *L*-Primordial-(Hhh)_2_ to 29 bp *D*-dsDNA and 29 bp *L-* dsDNA was assayed by surface plasmon resonance (SPR). We have previously shown that heterologously expressed *L*-Primordial-(HhH)_2_ binds to *D*-dsDNA by solution-state NMR and ELISA,^13^ and SPR experiments confirm that chemically synthesized *L*-Primordial-(HhH)_2_ also binds to *D*-dsDNA (**Figure 3C** and **D**). Binding of the *D*-protein to *L*-dsDNA corresponds to a *mirror world;* as such, the binding affinity of these two pairs should, in principle, be identical. The resulting binding curves are consistent with this expectation, with the *D*-protein:*L*-dsDNA pair having slightly lower affinity, likely due to some combination of inaccuracy in the protein concentrations and the lower synthetic purity of *D*-proteins and *L*-DNA relative to their natural counterparts^38, 39^.

Remarkably, binding of *D*-Primordial-(HhH)_2_ to *D*-dsDNA and *L*-Primordial-(HhH)_2_ to *L*-dsDNA was also observed (**Figure 3C** and **D**). The weaker binding of the *D*:*D* pair and the *L*:*L* pair relative to the *L*:*D* and *D*:*L* pairs is consistent with our expectation that, after nearly 4 billion years of coevolution, there should be significant optimization for the *L*:*D* chiral pair (and consequently for the *D*:*L* chiral pair). However, Ancestor-(HhH)_2_ (**Table 1**) – which is the direct result of ancestor sequence reconstruction and is comprised of a more complex alphabet of 15 different amino acid types, including aromatics – also binds to both *L*-dsDNA and *D*-dsDNA, and with an even smaller difference in affinity (**Figure 3D**). This observation confirms that functional ambidexterity is characteristic of the protein fold itself, and not unique to highly sequence-simplified variants. Dissociation kinetics of the natural chiral pair are complex and multiphasic and the kinetic constants associated with the mirror image pair are highly similar (**Figure S20**). Surprisingly, the dissociation kinetics of the *L*-protein:*L*-dsDNA pair are also highly similar to that of the natural and mirror image pairs (**Figure S21**), suggesting some degree of binding mode conservation.

To further confirm that binding is mediated by the PGIGP binding loop and the flanking □-helices, we mutated the PGIGP loop to GGGGG, yielding the construct *L*-Primordial-(HhH)_2_-5G (**Table 1**). This mutation does not change the charge of the protein nor does it occlude binding by insertion of a bulky residue. Instead, this mutation should increase the overall flexibility of the binding loops and the protein itself, and thus disfavor binding if a near-native conformation mediates the interaction with dsDNA. Although not as well-folded as the parent protein (**Figure 3B**), *L*-Primordial-(HhH)_2_-5G retains some □-helical character. As can been seen in **Figure 3D** and **E**, *L*-Primordial-(HhH)_2_-5G binds to *D*-dsDNA and *L*-dsDNA with drastically lower affinity than any other construct tested. These results confirm that binding is not simply mediated by promiscuous charge-charge interactions. The observations presented in **Figure 3** were confirmed on a second SPR chip (**Figure S22**) and are in agreement with microscale thermophoresis (MST) experiments (**Figure S23**), though MST experiments were concentration-limited due to significant changes in the initial fluorescence at high concentrations of DNA.

### The (HhH)_2_-Binding Mode is Largely Retained Upon Chiral Inversion

To probe the extent to which the binding mode between the natural and unnatural pairs resemble each other, MD simulations of *L*- and *D*-Primordial-(HhH)_2_ binding to *D*-dsDNA were performed. The Lenard-Jones short-range energies after 500 ns equilibration echoed experimental binding data, with binding of *L*-Primordial-(HhH)_2_ being the most stable and binding of *L*-Primordial-(HhH)_2_-5G being the weakest (**Figure S24**). *L*-Primordial-(HhH)_2_-5G dissociation from *D*-dsDNA was observed in one of the three simulations, also consistent with this being the least stable complex.

Bound structures after equilibration were clustered with a 2 Å cutoff and the largest cluster was used for further analysis (**Figure S25**). Heatmaps of the protein-dsDNA distances calculated from the largest cluster are shown is **Figure 4A**. For the natural, *L*-protein:*D*-dsDNA binding mode, four interaction surfaces are apparent. These surfaces correspond to the first and second PGIGP motifs of the (HhH)_2_-Fold (protein residues 14-18 and 46-50) binding to nucleotides of the ‘upper’ and ‘lower’ strands of the minor groove. Comparison to the *D*-protein:*D*-dsDNA binding mode reveals a remarkable conservation of the location of the interaction surfaces in three out of four cases (**Figure 4A**). As in the natural binding mode, surfaces are centered on the PGIGP motif, though with a slight C-terminal shift towards a trailing Arg residue. Therefore, the ambidextrous nature of the (HhH)_2_-Fold is not likely the result of a second, unique binding site. Heatmaps of *L*-Primordial-(HhH)_2_-5G binding to *D*-dsDNA show the greatest degree of distortion (**Figure S26**), with only one of the four binding surfaces relatively unperturbed (corresponding to the lower right interaction surface in the heatmap). Finally, inspection of a representative conformation taken from the largest binding cluster (**Figure 4B**) reveals that many of the features of the natural binding mode are similar in the binding of *D*-Primordial-(HhH)_2_ to *D*-dsDNA; namely, the participation of the N-terminus of the α-helix to binding the phosphodiester backbone of dsDNA, the partial insertion of the PGIG motif within the minor groove, and the binding of flanking Arg residues to nearby phosphates (see **SI** for selected movies).

**Figure 4.**
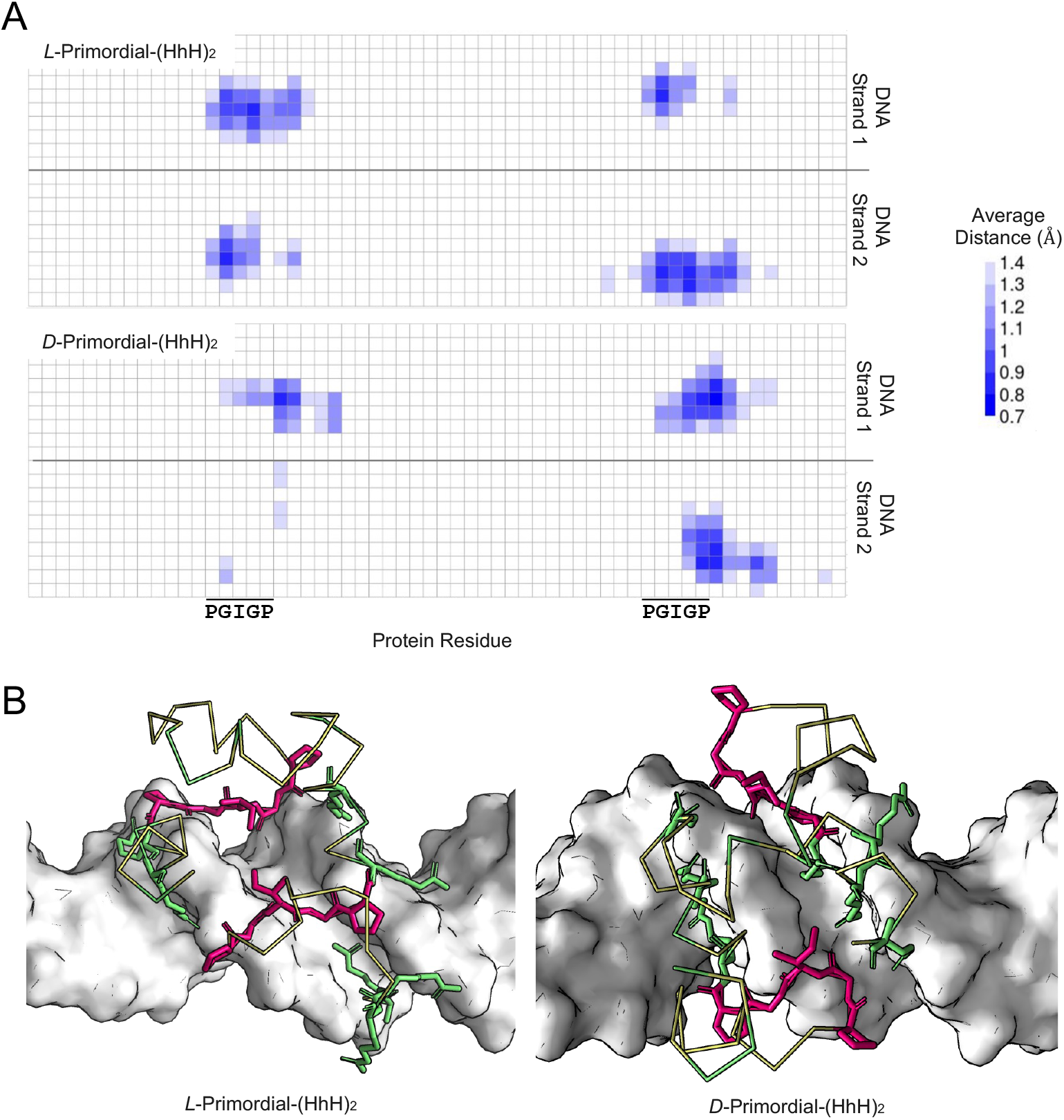
Molecular dynamics simulations of *L*- and *D*-Primordial-(HhH)2 bound to *D*-dsDNA. **A**. Heatmaps showing the minimum Cα to C/N distances between protein and DNA, respective, for the dominant binding clusters. Significant retention of the key binding surfaces upon chiral inversion is observed. **B**. Representative structures from the dominant binding clusters. PGIGP motifs are colored magenta and Arg residues are colored green. Backbone traces for the remainder of the protein is rendered as a yellow ribbon. Note that the PGIGP motif straddles the minor groove in both structures.

## Discussion

Total chiral inversion was tolerated along the entire putative evolutionary trajectory of the HhH protein family – from coacervation of a single HhH peptide with RNA to binding of the (HhH)_2_-Fold to dsDNA. These results suggest that functional ambidexterity is an enduring feature of this fold. Previously, GroEL/ES was determined to be functionally ambidextrous with respect to the protein substrate^40^, as it was able to refold DapA of either chirality into an active conformation. The GroEL/ES result can, and perhaps should, be rationalized as a consequence of the non-specific interactions made by chaperones and their need to accommodate diverse protein substrates. Similarly, the dimerization of transmembrane α-helices with inverted chirality was taken to mean that the interactions involved in dimerization were largely mediated by sidechains^41^. The case for the HhH protein family, however, is harder to interpret. First, with respect to coacervation, relatively little is known about the effects of chiral inversion and one of the few systems studied from this perspective exhibited dramatic effects^26^. Second, with respect to dsDNA binding, the interactions are specific, partially mediated by the backbone, and the same highly conserved residues participate in dsDNA binding of the natural and unnatural chiral pairs.

At present, it is not yet possible to say whether the functional ambidexterity of the HhH motif and the (HhH)_2_-Fold is a chance occurrence or a manifestation of early functional constraints. The pseudo-*C_2_* symmetry of the (HhH)_2_-Fold is certainly related to its functional ambidexterity, as it enables the approximately correct placement of the 4 α-helices even after chiral inversion. Symmetric proteins may have been preferred early in protein evolution, however, for reasons unrelated to functional ambidexterity; namely, because they can be realized by a minimal oligomerizing peptide^14, 17, 42, 43, 44, 45^. Moreover, in a world where only one chiral form of dsDNA and protein exists, there is little pressure to evolve specificity against mirror-image biopolymers. *If* the ambidexterity of this protein family is evolutionarily significant, it transforms our understanding of the history of homochirality, as ambidextrous domains would have unique properties in a complex ecosystem where both chiral forms of dsDNA or protein are in competition. Ultimately, for questions of this nature to be answered, other ancient protein folds will have to be resurrected and their functional ambidexterity assayed experimentally, a goal that is increasingly within reach^46^.

## Conclusions

The HhH motif and (HhH)_2_-Fold are ancient and ubiquitous nucleic acid binders. In an effort to better understand the history of homochirality on Earth, we now report that both the HhH motif and the (HhH)_2_-Fold exhibit signatures of functional ambidexterity. Whether this is coincidence or whether the history of homochirality is written into the most ancient protein folds remains to be seen. If the latter case is true, it suggests that competition and biological selection – not abiotic chemical or physical processes – could have been the ultimate decider of the chiral preferences of life on Earth. Regardless, the HhH protein family is an intriguing exception to the dogma of reciprocal chiral substrate specificity proposed by Milton and Kent. The veil between worlds with alternative chiral preferences may not be as impenetrable as is often assumed.

## Acknowledgements

We acknowledge Dr. Yosef Scolnik for assistance collecting circular dichroism spectra.

N.M. acknowledges financial support from the Israel Science Foundation (1388/22). O.W.-K. is supported by the Kaete Klausner Fellowship.

