## Supplemental Information for "Functional Ambidexterity of an Ancient Nucleic Acid-Binding Domain"

**Supplemental Materials and Methods**

**Reagent sourcing**. Buffers were prepared using MilliQ water (Millipore, Merck). Ultrapure guanidinium chloride (Gn·HCl, Chem-Impex Inc) was used in all ligation reactions. $Na_2HPO_4 \cdot 12H_2O$, ethanedithiol (EDT), triisopropylsilane (TIPS), 4-mercaptophenylacetic acid (MPAA) were purchased from Sigma-Aldrich (Rehovot, Israel). Cy5-NHS ester was purchased from BroadPharm® (USA). Acetylacetone and tris(2-carboxyethyl)phosphine (TCEP) were purchased from Tokyo Chemical Industry. *p*-nitrophenyl chlorophormate was purchased from Acros Organics. All Fmoc-amino acids (*D*- and *L*-) were obtained from CS Bio Co. (Menlo Park, CA), Matrix Innovation (Quebec City, Canada) or Chem-Impex Inc., with the following side chain protecting groups: Arg(Pbf), Glu(OtBu), Gly(OtBu), Ser(tBu), Thr(tBu), (Pbf = 2,2,4,6,7-pentamethyl-2,3-dihydrobenzofuran-5-sulfonyl). TentaGel® R RAM resin (loading 0.19 mmol/g) was purchased from Rapp Polymer GmbH (Germany) and 2-chlorotrityl-resin (loading of 0.3-0.8 eq/g) was purchased from Chem-Impex Inc. 1-[Bis(dimethylamino)methylen]-5-chlorobenzotriazolium 3-oxide hexafluorophosphate and *N,N,N',N'*-Tetramethyl-O-(6-chloro-1H-benzotriazol-1-yl)uronium hexafluorophosphate (HCTU) was purchased from Luxembourg Biotechnologies Ltd. (Rehovot, Israel). All solvents: *N,N*-dimethylformamide (DMF), dichloromethane (DCM), acetonitrile (ACN), *N,N*-diisopropylethyl amine (DIEA), Trifluoroacetic acid (TFA), piperidine (Pip), dimethylsulfoxide (DMSO) and Boc-anhydride were purchased from Bio-Lab (Jerusalem, Israel) and were peptide synthesis, HPLC or ULC-grade.

**High Performance Liquid Chromatography (HPLC)**. Analytical reversed-phase HPLC (RP-HPLC) was performed on a Waters Alliance HPLC with 220 and 280 nm UV detection using an XBridge BEH300 C4 column (3.5 µm, 130 Å, 4.6 × 150 mm). Semi-preparative RP-HPLC was performed on a XBridge BEH C4 column (5 µm, 300 Å, 10 × 150 mm) and XSelect CSH C18 column (5 µm, 130 Å, 10 × 150 mm). Preparative RP-HPLC was performed on a XSelect C4 column (5 µm, 130 Å, 19× 250 mm) or XSelect C18 column (5 µm, 30 × 250 mm). The flow rates were 1 mL/min (analytical), 3.35 mL/min (Semi-preparative), or 10-20 mL/min (preparative). Linear gradients of ACN (with 0.1% TFA, eluent B) in water (with 0.1 % TFA, eluent A) were used for all systems to elute bound peptides.

**Electrospray Ionization Mass Spectrometry (ESI-MS).** ESI-MS was performed on LCQ Fleet Ion Trap mass spectrometer (Thermo Scientific). Peptide masses were calculated from the experimental mass to charge (m/z) ratios from the observed multiply charged species of a peptide. Deconvolution of the experimental MS data was performed with MagTran v1.03.

**High-Resolution Mass Spectrometry (HR-MS)**

HR-MS spectra were recorded on a Q Exactive Plus Orbitrap mass spectrometer (Thermo Scientific) with an ESI source and 140,000 FWHM, in a method with the automatic gain control (AGC) target set to 1E6 and a scan range of 400-2800 m/z. Deconvolution of the raw MS data was performed with MagTran v1.03.

**Peptide Synthesis**

**General procedure for Fmoc-Solid Phase Peptide Synthesis (SPPS)**: Peptides were prepared by automatic peptide synthesizer (CS136XT, CS Bio Inc. CA) typically on 0.25 mmol scales. Fmoc-protected amino acids (2 mmol in 5 mL DMF) were activated with HCTU (2 mmol in 5 mL DMF) and DIEA (4 mmol in 5 mL DMF) for 5 min and allowed to couple for 25 min, with constant shaking. Fmoc-deprotection was carried out with 20% piperidine in DMF (2 × 5 min).

**Cleavage, deprotection and purification**: The peptide-resins were washed with DMF and DCM and then dried under vacuum. The dried peptide-resins were deprotected and simultaneously cleaved using a TFA/water/thioanisole/triisopropylsilane/ethanedithiol (92.5:1.5:1.5:1.5:1.5) cocktail for 4 h. The cleavage mixtures were filtered and TFA was evaporated with $N_2$-bubbling to a minimum volume, to which an eightfold volume of cold ether was added dropwise. The precipitated crude peptides were centrifuged (5000 rpm, 10 min), ether was removed, and the crude peptide was dissolved in ACN/water (1:1) containing 0.1% TFA and was further diluted to ca. 25% ACN with water and lyophilized.

**Synthesis of *L*-Primordial-(HhH)$_2$**

***Sequence:***

RIRRASVEELTEVPGIGPRLARRILER**<u>LA</u>**SIERIRRASVEELTEVPGIGPRLARRILERL

*L*-Primordial-(HhH)$_2$ was prepared from two peptide segments using the native chemical ligation (NCL) and desulfurization approach (**Scheme S1**). The two peptide segments were the thioester surrogate *L*-Primordial(1-28)-Nbz (Nbz = *N*-acylurea) (Blanco-Canosa & Dawson, 2008), and *L*-Primordial (29-60)(A29C), in which Ala29 was temporary substituted with Cys to allow for the NCL reaction, and will later be desulfurized to natural Ala29 post-ligation. The ligation site within the sequence (above) is shown in bold and underlined.

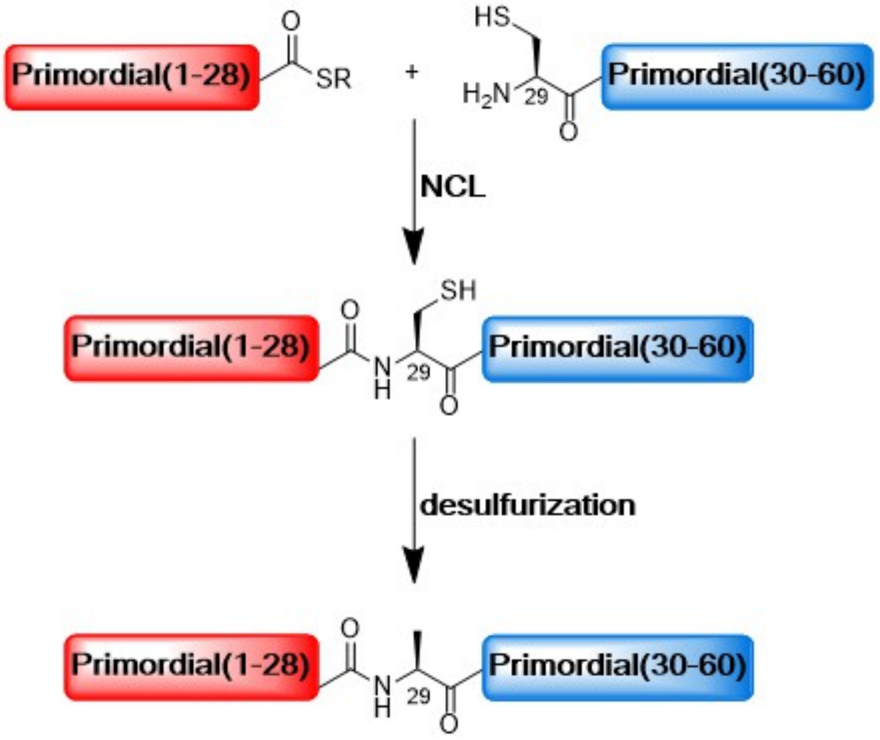


**Scheme S1**. **Synthesis of *L*-Primordial-(HhH)$_2$**. Chemical protein synthesis scheme for *L*-Primordial-(HhH)$_2$. The protein was synthesized from two half-peptides and then joined using NCL and desulfurization. The N-terminal half-peptide bears a C-terminal thioester surrogate (shown here as thioester for simplicity), and the C-terminal peptide bears an N-terminal cysteine residue. After peptide ligation, the cysteine residue is desulfurized to yield alanine.

***Synthesis of L-Primordial(1-28)-Nbz peptide:***

*L*-Primordial(1-28)-Nbz was synthesized first on Fmoc-Dbz-resin (0.25 mmol scale) with an automated peptide synthesizer. Mono-Fmoc-3,4-diaminobenzoic acid (Fmoc-Dbz-OH, 3 equiv) (Blanco-Canosa & Dawson, 2008) was activated with HCTU (3 equiv)/DIEA (6 equiv) in DMF and was doubly coupled to the free amine of TentaGel® R RAM resin (0.19 mmol/g, 0.25 mmol scale) for 1 h. The first amino acid, Leu28, was also doubly coupled.

*N-terminal Boc-protection:* After synthesis completion, the Fmoc protecting group of the N-terminal Arg was removed and the peptide-resin was treated with 1.5 equiv of Boc-anhydride solution dissolved in 10 mL DCM and 2.0 equiv DIEA. The reaction was left overnight to give N-terminal Boc-protected peptide-resin required before the step of Dbz to Nbz conversion.

*On-resin Nbz formation:* The resin was washed with DCM and a solution of *p*-nitrophenyl chloroformate (5 equiv, 1.25 mmol) in DCM (5 mL) was added, shaken for 1 h at 25 °C and washed with DCM (3 × 5 mL) and DMF (3 × 5 mL). This step was repeated one more time. Following this, the resin was washed with DMF and a 5 mL solution of 0.5 M DIEA in DMF was added and shaken for an additional 30 min to complete the cyclization/Nbz formation (repeated twice), and washed with DMF (3 × 5 mL) and DCM (3 × 5 mL) and dried under vacuum.

*Deprotection and cleavage:* The peptide-resin was deprotected and cleaved as described previously to give 909 mg of crude *L*-Primordial(1-28)-Nbz.

*Purification of L-Primordial(1-28)-Nbz:* 150 mg of crude peptide were taken and dissolved in 25% ACN with water and purified by preparative RP-HPLC (XSelect C18 column, 5 µm, 130 Å, 30 × 250 mm), using a gradient of 30-60% B over 42 min to give pure *L*-Primordial(1-28)-Nbz (21 mg, 15% yield). The HPLC analysis (**Figure S1**) was carried out on a C4 analytical column (XBridge BEH300, 3.5 µm, 130 Å, 4.6 × 150 mm) using a gradient of 5% B over 2 min then 5-70% B over 20 min).

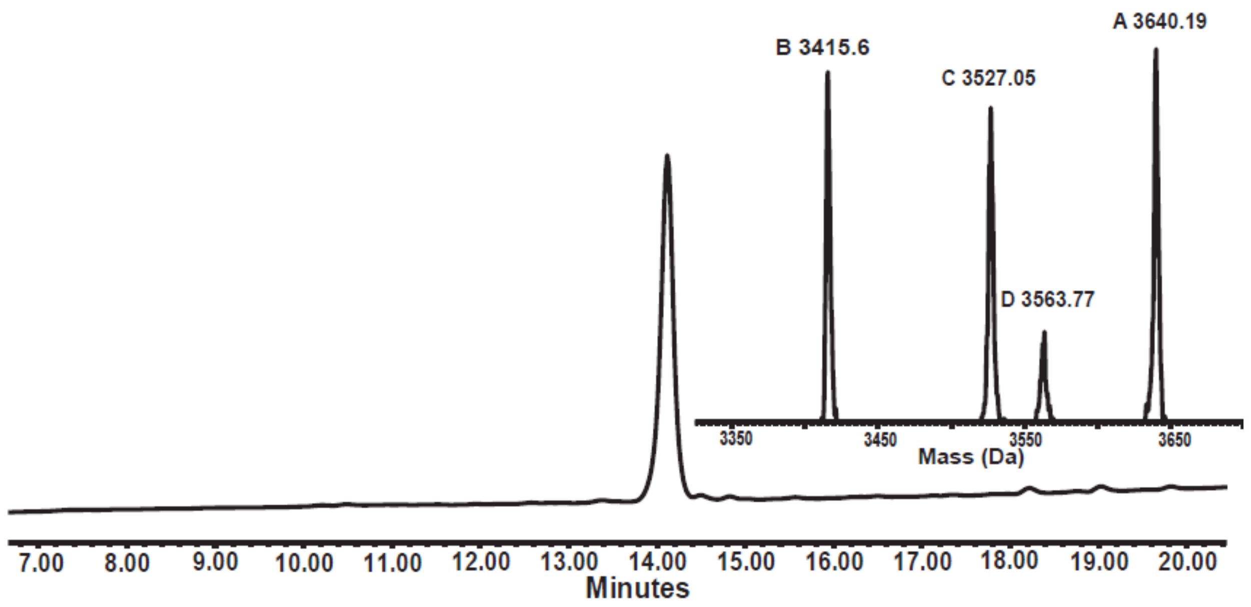


**Figure S1**. HPLC chromatograms and ESI-MS for purified *L*-Primordial(1-28)-Nbz, with the inset showing the corresponding mass (calc. 3416.0 Da; obs. 3415.6 Da, also observed [M+1 TFA] 3527.05 Da, [M+2 TFA] 3640.19 Da; the 3563.77 Da species is an impurity).

***Synthesis of the second segment, L-Primordial(29-60)(A29C):***

The peptide was synthesized on 2-chlorotrityl-resin (loading 0.3 mmol/g, 0.25 mmol scale) on an automated peptide synthesizer. Ala29 was substituted with Cys to permit Cys-NCL with the first segment *L*-Primordial(1-28)-Nbz.

*Deprotection and cleavage:* Peptide was deprotected and cleaved as described previously to give 1356 mg of crude *L*-Primordial(29-60)(A29C).

*Purification:* The peptide was purified by RP-HPLC (200 mg of crude) (XSelect C18 column, 5 µm, 130 Å, 30 × 250 mm) using a gradient of 30-60% B over 42 min to give pure Primordial(29-60)(A29C) (68 mg, 34% yield). The HPLC analysis (**Figure S2**) was carried out on a C4 analytical column.

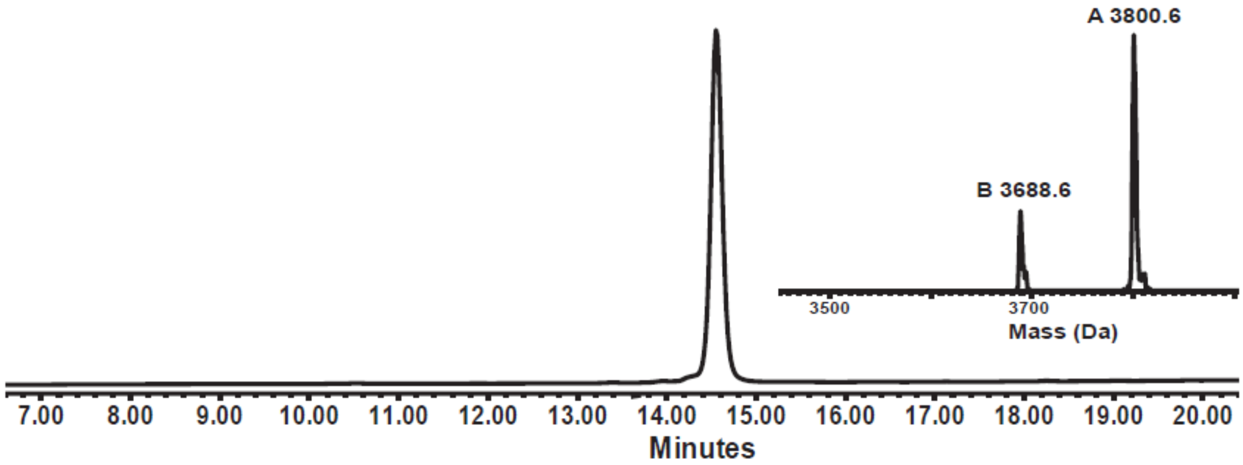


**Figure S2**. HPLC chromatograms and ESI-MS for purified *L*-Primordial(29-60)(A29C), with the inset showing the corresponding mass (calc. 3689.3 Da; obs. 3688.6 Da, [M+TFA] 3800.6 Da).

*NCL between L-Primordial(1-28)-Nbz and L-Primordial(29-60)(A29C):* For ligation, *L*-Primordial(1-28)-Nbz peptide (~10.5 mg, 3.1 µmol, final conc. ~1 mM) was dissolved in 3 mL of argon-degassed phosphate buffer (200 mM $NaH_2PO_4$, 6 M Gn·HCl, 0.2 M MPAA, 0.05 M TCEP , pH 6.5) and this mixture was added to *L*-Primordial(29-60)(A29C) peptide (~16 mg, 4.34 µmol, final conc. ~1.3 mM). The progress of the reaction was followed by analytical HPLC (XBridge C4 column, 3.5 µm, 4.6 × 150 mm) with a gradient of 10-60% B over 20 min. The ligation was quenched after 4 h. The ligation product was purified by semi-preparative HPLC (XBridge BEH300 C4 column, 5 µm, 19 × 150 mm, method 25-50% B over 45 min) to yeild the corresponding *L*-Primordial(1-60)(A29C)-$(HhH)_2$ product (10 mg, 38% yield; **Figure S3**).

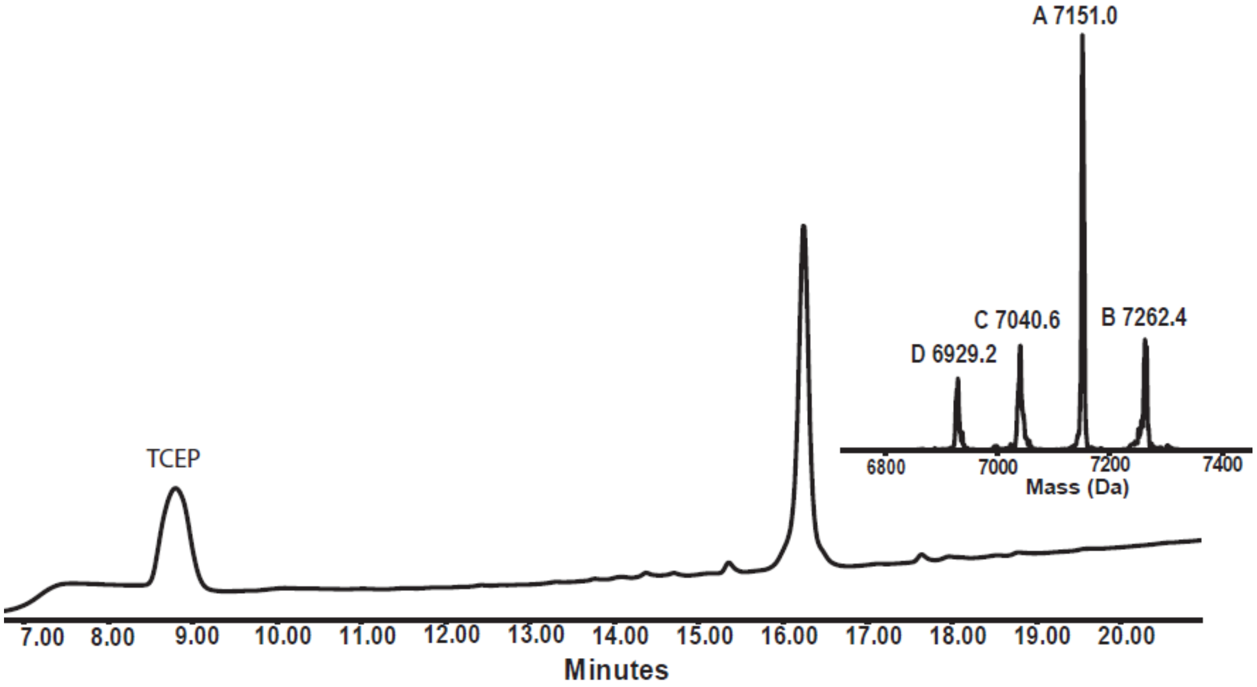


**Figure S3**. HPLC chromatograms and ESI-MS for purified *L*-Primordial-(1-60)(A29C)-$(HhH)_2$, with the inset showing the corresponding mass (calc. 6928.20 Da; obs 6929.2 Da, [M+TFA] 7040.6 Da, [M+2TFA] 7151.0 Da, [M+3TFA] 7262.4 Da).

*Desulfurization of L-Primordial(1-60)(A29C)-$(HhH)_2$ to give L-Primordial-$(HhH)_2$:* L-Primordial(1-60)(A29C)-$(HhH)_2$ (17 mg, 2.46 μmol) was dissolved in 4 mL of argon degassed phosphate buffer (200 mM $NaH_2PO_4$, 6 M Gn·HCl, 400 equiv TCEP, 200 equiv VA-044, pH 5.5), and left for 48 hours to afford the desired product. The progress of the reaction (**Figure S4**) was followed by analytical HPLC (XBridge C4 column, 3.5 μm, 4.6 × 150 mm) with gradient 10-60% B over 20 min. *L*-Primordial-$(HhH)_2$ was purified by semi-preparative HPLC (XBridge BEH300 C4 column, 5 μm, 19 × 150 mm, method 25-50% B over 45 min) (8 mg, 47% yield).

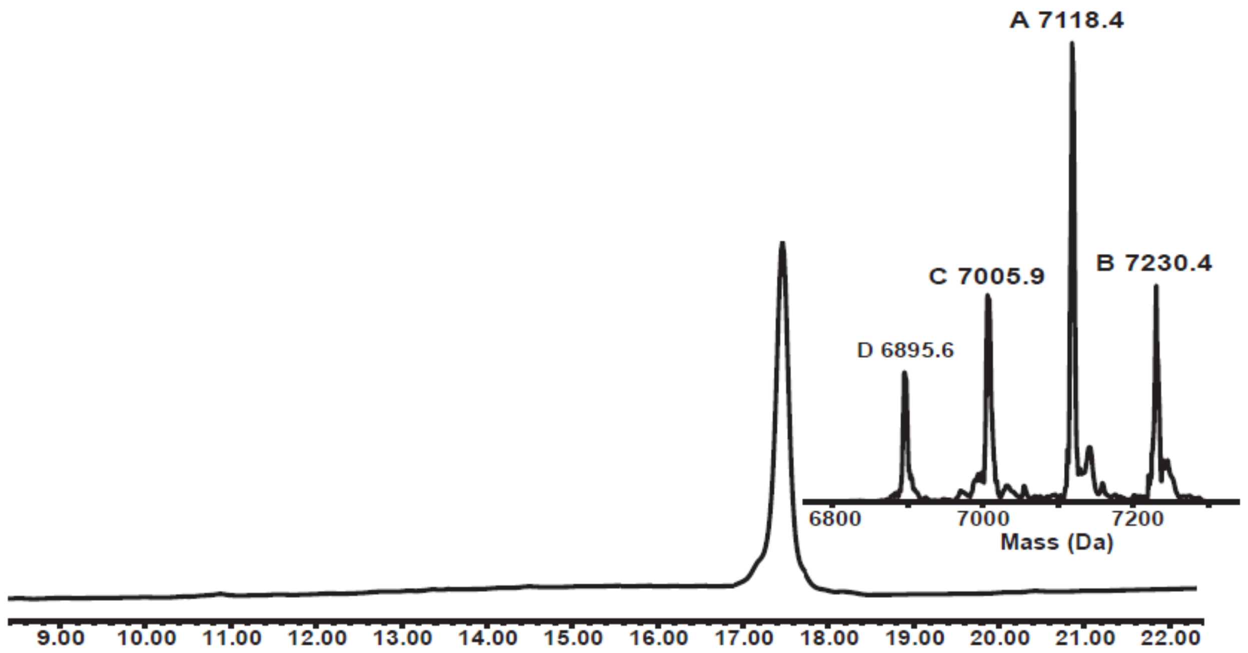


**Figure S4**. Desulfurization of *L*-Primordial-(1-60)(A29C)-$(HhH)_2$. Analytical HPLC of the desulfurization reaction after 48 h with the ESI-MS of desired product *L*-Primordial-$(HhH)_2$ (calc. 6896.1 Da; obs. 6895.6 Da, [M+TFA] 7005.9 Da, [M+2 TFA] 7118.4 Da, [M+3 TFA] 7230.4 Da).

**Synthesis of *D*-Primordial-$(HhH)_2$**

***Sequence (same as the L-Primordial-$(HhH)_2$)*:**

d(RIRRASVEELTEVPGIGPRLARRILER**LA**SIERIRRASVEELTEVPGIGPRLARRILERL)

The *D*-form of the Primordial-$(HhH)_2$ protein was prepared using the same approach as for the *L*-protein, from two peptide segments using NCL and desulfurization (**Scheme S1**). The two peptide segments were *D*-Primordial(1-28)-$NHNH_2$, and *D*-Primordial(29-60)(A29C), in which Ala29 was temporary substituted with Cys to allow for the NCL reaction, and was then desulfurized to natural Ala29 after ligation. The ligation site within the sequence (above) is shown in bold and underlined.

***Synthesis of D-Primordial(1-28)-COSR peptide:***

For the synthesis of *D*-Primordial(1-28)-COSR, we first prepared the C-terminal hydrazide thioester surrogate, which was carried out on 2-chlorotrityl chloride-resin. First, the resin (0.25

mmol scale; loading 0.5 mmol/g) was swelled in DMF for 1 h and treated twice with freshly prepared 5% hydrazine in DMF for 1 h and decanted (Zheng et al., 2013). The resin was washed well with DMF and then treated with 10% MeOH in DMF for 30 min. The hydrazine functionalized chlorotrityl-resin was used for standard Fmoc-SPPS, with the coupling of the *D*-amino acids performed using automated synthesizer, while the *D*-Ile residues were manually coupled.

*Deprotection and cleavage:* Peptide was deprotected and cleaved off the resin as described previously to give 400 mg of crude *D*-Primordial(1-28)-$NHNH_2$.

*Purification:* The peptide was purified by RP-HPLC (100 mg of crude peptide) on an XSelect C18 column (5 µm, 130 Å, 30 × 250 mm) using a gradient of 30-60% B over 42 min to give pure *D*-Primordial(1-28)-$NHNH_2$ (15 mg, 15% yield). The HPLC analysis (**Figure S5**) was carried out on a C4 analytical column.

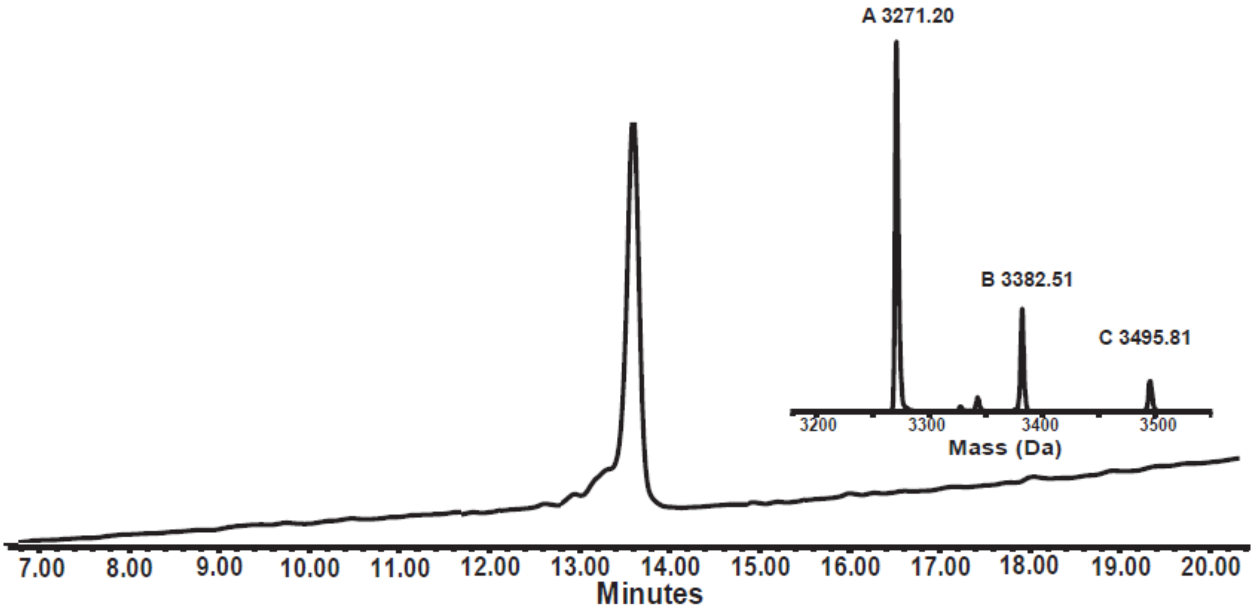


**Figure S5**. HPLC chromatograms and ESI-MS for purified *D*-Primordial(1-28)-$NHNH_2$, with the inset showing the corresponding mass (calc. 3270.89 Da; obs. 3271.20 Da, [M+TFA] 3382.51 Da, [M+2 TFA] 3495.81 Da).

*Procedure for thioesterification*

The conversion to thioester was done by dissolving 15 mg of peptide in phosphate buffer (200 mM, 6 M Gn·HCl, pH ~2.5; final conc. ~1 mM) and treated with 50 equivalents of acetylacetone (acac) and 200 equiv of MPAA for 2 h at room temperature (Flood et al., 2018). This reaction mixture was kept for the next ligation step.

***Synthesis of the second segment, D-Primordial(29-60)(A29C):***

This peptide was synthesized on an automated synthesizer with manual coupling of *D*-Ile residues and cleaved as described previously for *L*-Primordial(29-60)(A29C) to obtain 405 mg of crude *D*-Primordial(29-60)(A29C).

*Purification:* The peptide was purified by RP-HPLC (100 mg of crude) (XSelect C18 column, 5 µm, 130 Å, 30 × 250 mm) using a gradient of 30-60% B over 42 min to give pure *D*-Primordial(29-60)(A29C) (26 mg, 26% yield). The HPLC analysis (**Figure S6**) was carried out on a C4 analytical column.

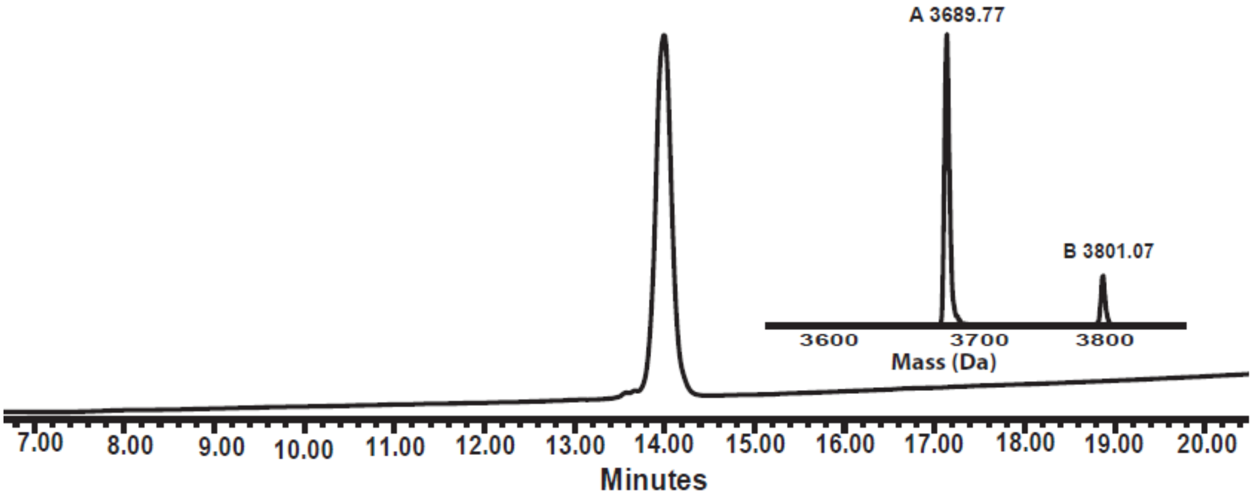


**Figure S6**. HPLC chromatograms and ESI-MS for purified *D*-Primordial(29-60)(A29C), with the inset showing the corresponding mass (calc. 3689.35 Da; obs. 3689.77 Da, [M+TFA] 3801.07 Da).

*NCL between D-Primordial(1-28)-COSR and D-Primordial(29-60)(A29C):* For the ligation reaction, 200 equivalents of TCEP were added in the thioesterification reaction mixture and the pH was adjusted to 6.5. Then, *D*-Primordial(29-60)(A29C) peptide (~22.5 mg, 6.1 µmol, final conc. ~1.3 mM) was added. The progress of the reaction was followed by analytical HPLC (XBridge C4 column, 3.5 µm, 4.6 × 150 mm) with gradient of 10-60% B over 20 min. The ligation was left overnight. Ligation product was purified by semi-preparative HPLC (XBridge BEH300 C4 column, 5 µm, 19 × 150 mm, method 25-50% B over 45 min) to afford the corresponding *D*-Primordial(1-60)(A29C)-$(HhH)_2$ (7 mg, 19% yield; **Figure S7**).

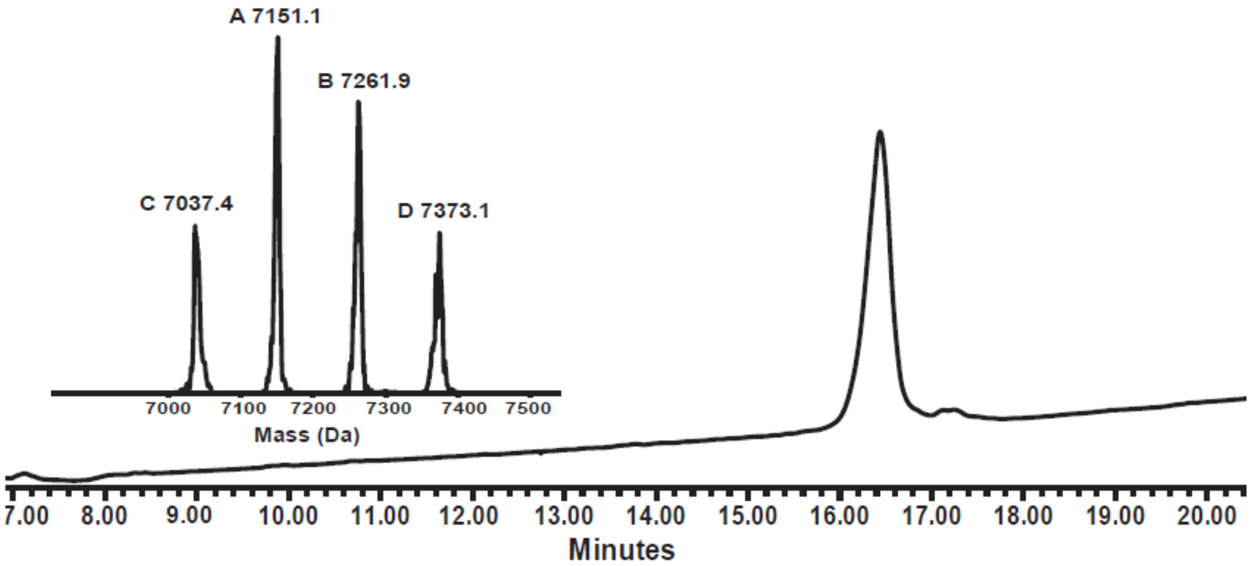


**Figure S7**. HPLC chromatograms and ESI-MS for purified *D*-Primordial(1-60)(A29C)-$(HhH)_2$, with the inset showing the corresponding mass (calc. 6928.2 Da, [M+TFA] 7042.2 Da; obs. [M+TFA] 7037.4 Da, [M+2 TFA] 7151.1 Da, [M+3 TFA] 7261.9 Da, [M+4 TFA] 7373.1 Da).

*Desulfurization of D-Primordial(1-60)(A29C)-$(HhH)_2$ to give D-Primordial-$(HhH)_2$:* The desulfurization reaction was performed as described previously on 7 mg of *D*-Primordial(1-60)(A29C)-$(HhH)_2$. The progress of the reaction was followed by analytical HPLC (XBridge C4 column, 3.5 µm, 4.6 × 150 mm) with a gradient of 10-60% B over 20 min. *D*-Primordial-$(HhH)_2$, was purified by semi-preparative HPLC (XBridge BEH300 C4 column, 5 µm, 19 × 150 mm, method 25-50% B over 45 min) (**Figure S8**) to obtain 3.5 mg of *D*-Primordial-$(HhH)_2$ (50 % yield).

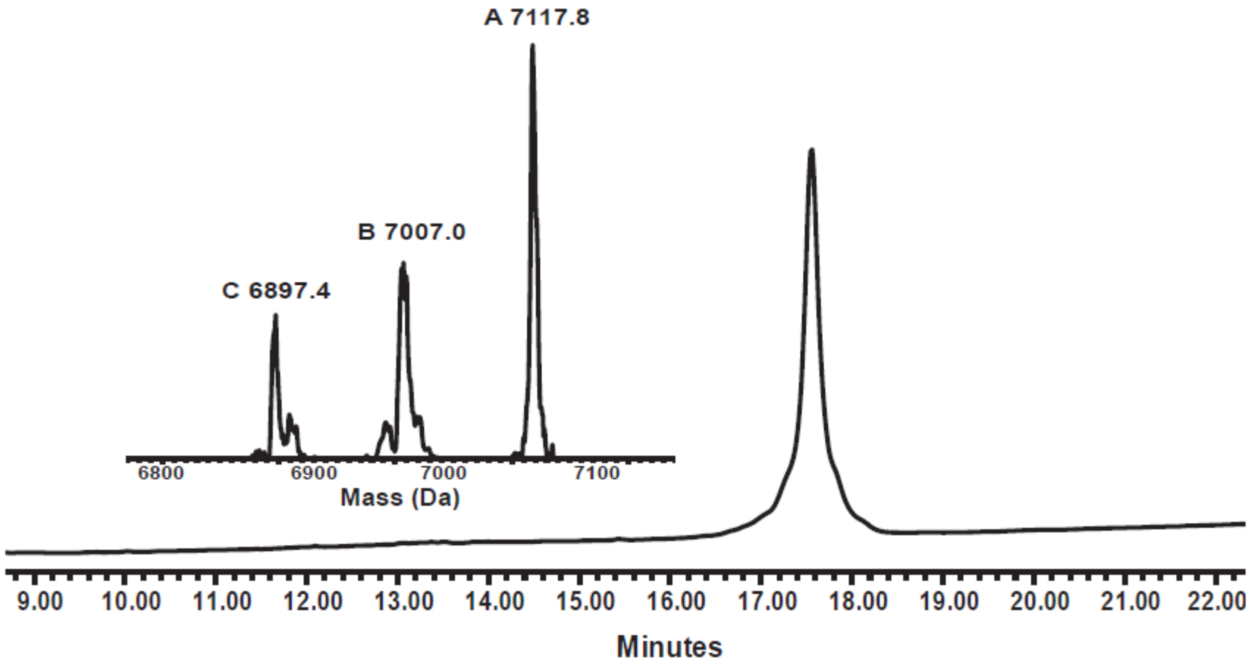


**Figure S8**. Desulfurization of *D*-Primordial(1-60)(A29C)-(HhH)$_2$. Analytical HPLC of the desulfurization reaction after 48 h with ESI-MS of desired product *D*-Primordial-(HhH)$_2$ (calc. 6896.1 Da; obs. 6897.4 Da, [M+TFA] 7007.0 Da, [M+ 2 TFA] 7117.8 Da).

**Synthesis of *L*-Primordial-(HhH)$_2$-5G**

***Sequence:***

RIRRASVEELTEV***GGGGG***RLARRILER**<u>LA</u>**SIERIRRASVEELTEV***GGGGG***RLARRILERL

The *L*-form of the disrupted Primordial protein was prepared from two peptide segments, NCL and desulfurization approach (**Scheme S2**). The sequence was similar to that of Primordial-(HhH)$_2$ except that the ***PGIGP*** binding motif was replaced by ***GGGGG*** (*bold and italics*). The two peptide segments were Primordial(1-28)-5G-$NHNH_2$, and Primordial(29-60)(A29C)-5G, in which Ala29 was temporary substituted with Cys to allow for NCL reaction, and was later desulfurized to natural Ala29 after ligation. The ligation site is shown in *bold* and *underlined*. HPLC chromatograms and ESI-MS for purified peptide segments and ligation product are presented in **Figures S9-S11**.

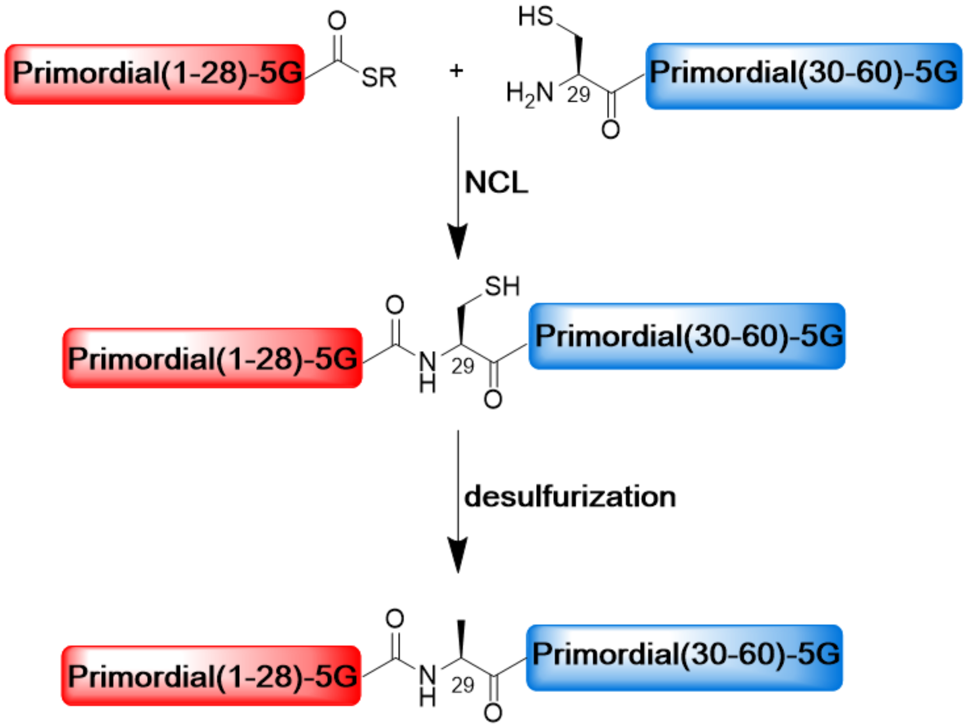


**Scheme S2**. **Synthesis of *L*-Primordial-(HhH)$_2$-5G**. Chemical protein synthesis scheme for *L*-Primordial-(HhH)$_2$-5G. The protein was prepared from two half-peptides and then joined using an NCL and desulfurization approach. The N-terminal half-peptide bears the C-terminal thioester, and the C-terminal peptide bears an N-terminal cysteine residue. After peptide ligation, the Cys residue is desulfurized to yield natural Ala.

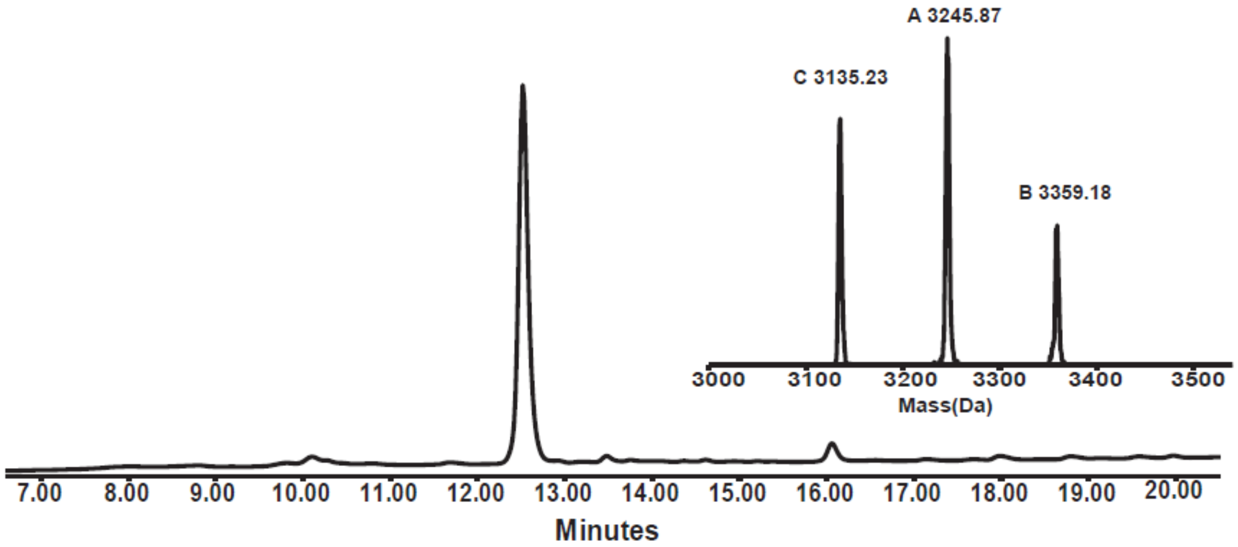


**Figure S9**. HPLC chromatograms and ESI-MS for purified *L*-Primordial(1-28)-5G-$NHNH_2$, with the inset showing the corresponding mass (calc. 3136.63 Da; obs. 3135.23 Da, [M+TFA] 3245.87 Da, [M+2 TFA] 3359.18 Da).

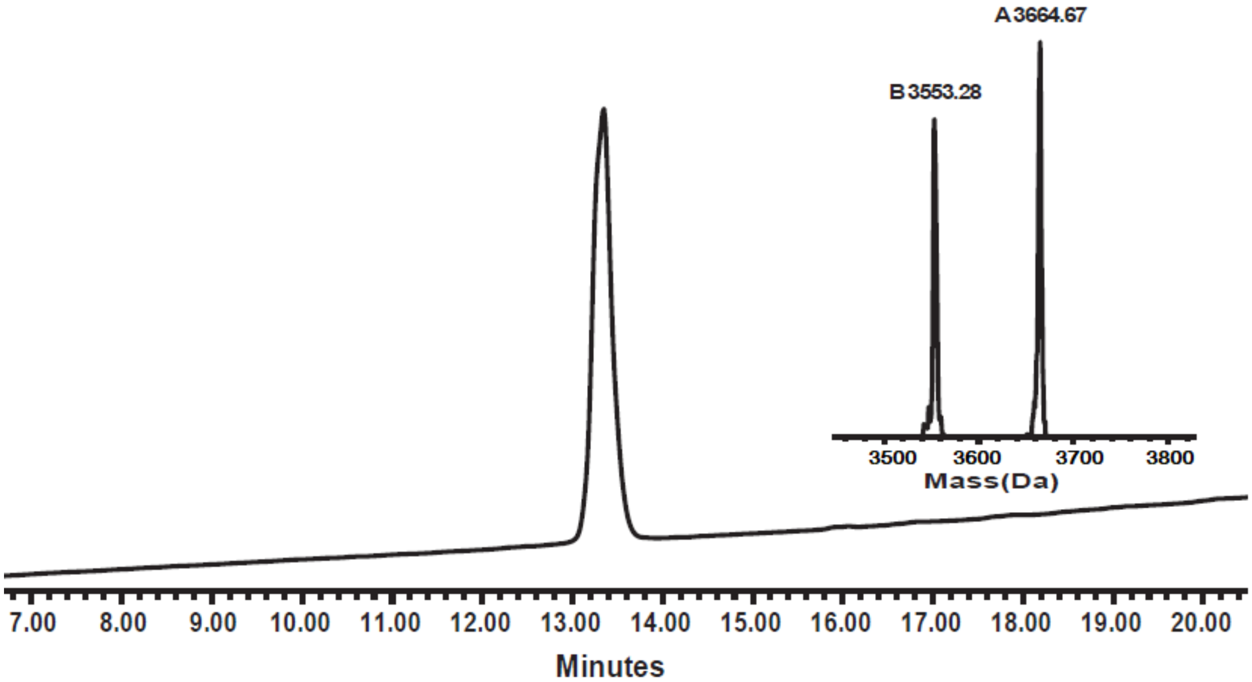


**Figure S10**. HPLC chromatograms and ESI-MS for purified *L*-Primordial(29-60)(A29C)-5G, with the inset showing the corresponding mass (calc. 3553.12 Da; obs. 3553.28 Da, [M+TFA] 3664.67 Da).

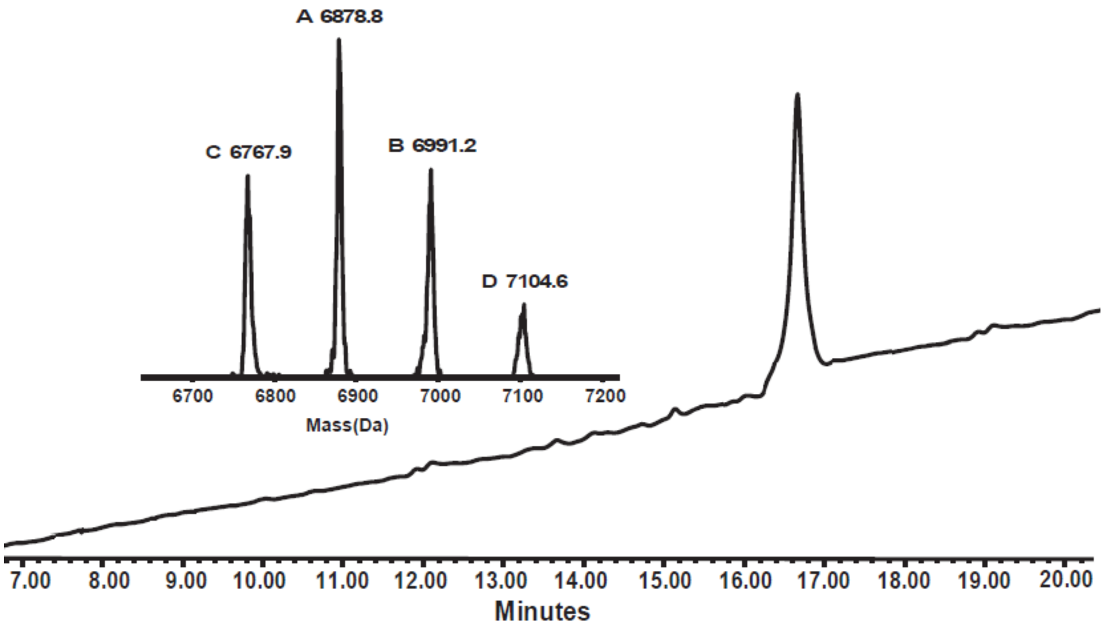


**Figure S11**. HPLC chromatograms and ESI-MS for *L*-Primordial-$(HhH)_2$(A29C)-5G with the inset showing the corresponding mass (calc. 6655.7 Da, [M+TFA] 6769.7 Da; obs. 6767.9 Da, [M+ 2 TFA] 6878.8 Da, [M+3 TFA] 6991.2 Da, [M+4 TFA] 7104.6 Da).

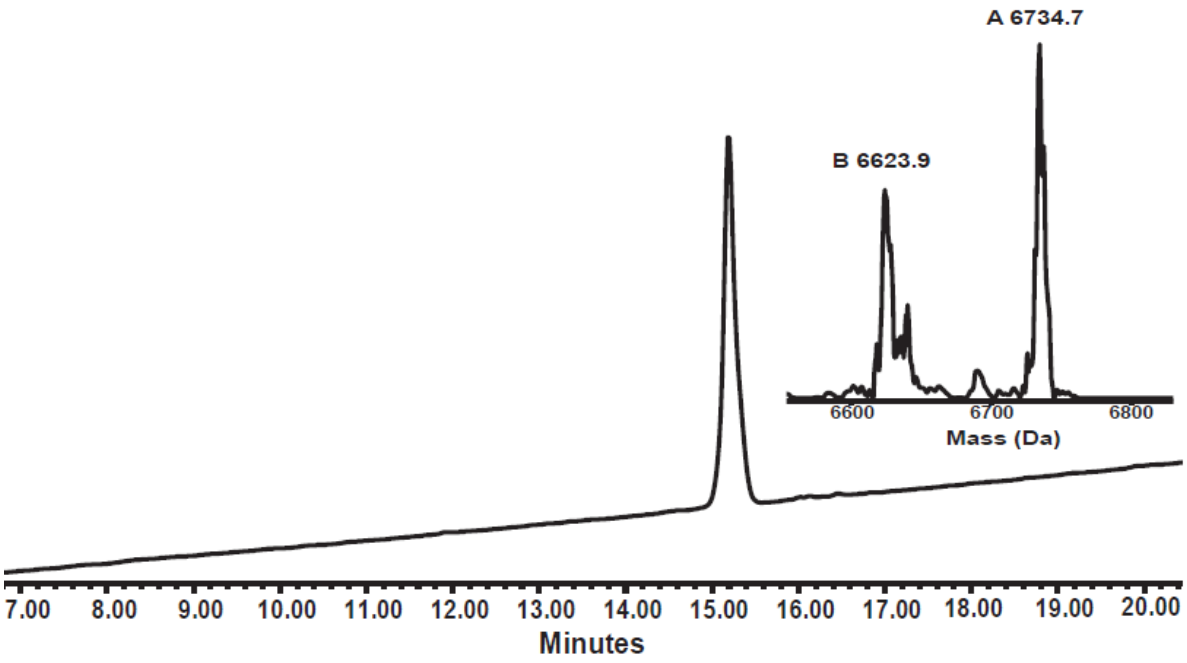


**Figure S12**. HPLC chromatograms and ESI-MS for purified *L*-Primordial-$(HhH)_2$-5G with the inset showing the corresponding mass (calc. 6623.7 Da; obs. 6623.9 Da, [M+ TFA] 6734.7 Da).

**Synthesis of Ancestor-$(HhH)_2$**

The total synthesis of Ancestor-$(HhH)_2$ is described in the **SI** of our previous article (Longo et al., 2020).

**Synthesis of D/L-Precursor-HhH**

***Sequence:***

RIRRASVEELTEVPGIGPRLARRILERLA

Single HhH peptides in which *D*- and *L*-amino acids were coupled alternatively (*D*-amino acids are underlined in the sequence shown above) were synthesized by an automatic peptide synthesizer (CS136XT, CS Bio Inc. CA) on a 0.15 mmol Rink amide resin (RAPP Polymer, loading 0.19) as described above. Arg residues were doubly coupled and all the *D*-amino acids were manually coupled. The peptide *D/L*-Precursor-HhH was then cleaved as described above yielding 264 mg of crude peptide.

*Purification:* The peptide was purified by RP-HPLC (50 mg of crude peptide) on an XSelect C4 column (5 µm, 130 Å, 19× 250 mm) using a gradient of 25-45% B over 42 min to give pure *D/L*-Precursor-HhH (8 mg, 16% yield). The HPLC analysis (**Figure S13**) was carried out on a C4 analytical column.

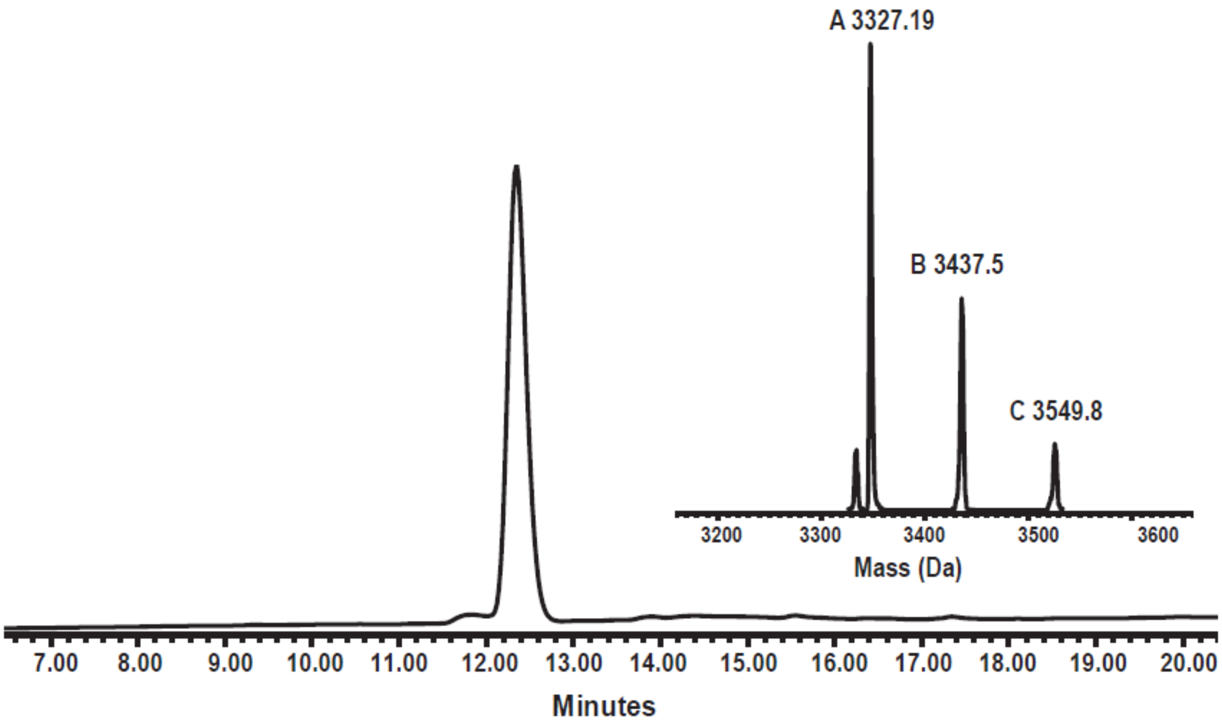


**Figure S13**. HPLC chromatograms and ESI-MS for purified *D/L*-Precursor-HhH with the inset showing the corresponding mass. (calc. 3326.96 Da; obs. 3327.19 Da, [M+TFA] 3437,5 Da, [M+2 TFA] 3549.8 Da).

**Proteins labeled with Cy5-NHS Ester for MST studies**

*L*-Primordial-$(HhH)_2$ and *L*-Primordial-$(HhH)_2$-5G were labeled with Cy5-NHS Ester for measurement with Microscale Thermophoresis (MST). For the labeling reaction, a stock solution of labeling reagent was prepared by dissolving 1 mg of Cy5-NHS ester in 200 µL DMF (8.12 mM). Then, 1 mg of protein was dissolved in 0.5 mL 0.2 M PBS pH 8.5 (conc. ~0.29 mM), and 71 µL of the Cy5-NHS ester stock solution (4 equiv, conc. ~1.16 mM) were added to the protein and vortexed well. The reaction was kept at room temperature for approximately 6 hours [https://www.lumiprobe.com/protocols/nhs-ester-labeling]. The reaction was monitored by HPLC and ESI-MS. Labeled peptides were purified by an XSelect CSH C18 column (5 µm, 130 Å, 10 × 150 mm) using a gradient of 30-50% B over 55 min. The labeled peptides were further analyzed using HPLC on a C4 analytical column (**Figures S14-S15**). Yield of the labeling reactions were 70-85% after the purification step.

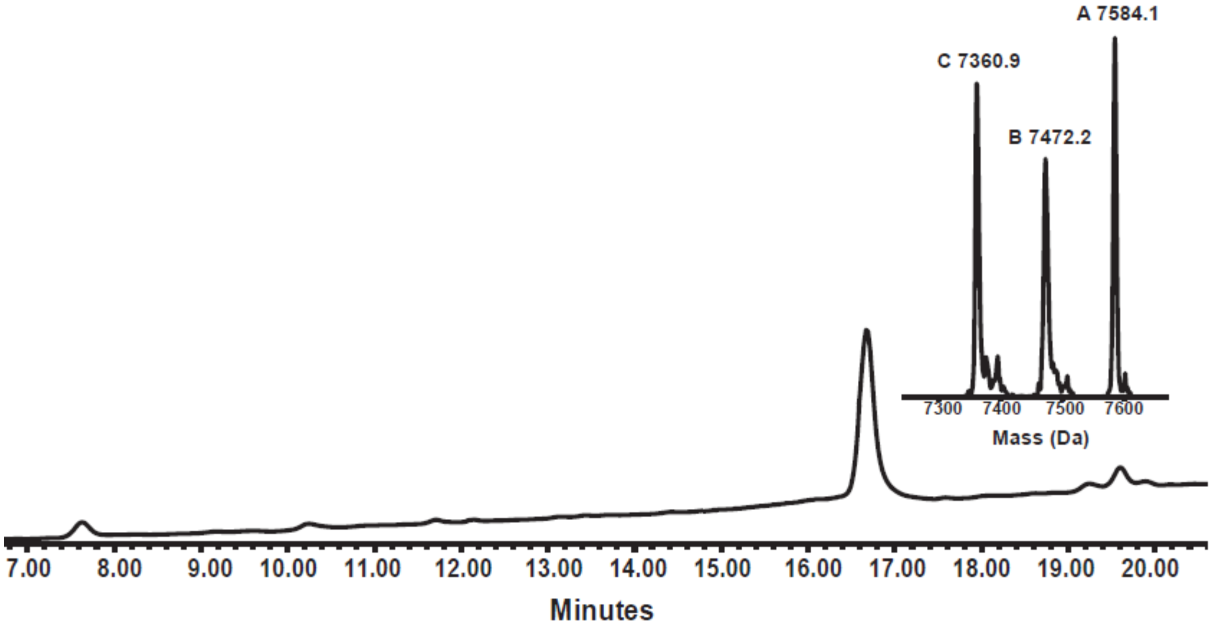


**Figure S14**. HPLC chromatograms and ESI-MS for purified *L*-Primordial-$(HhH)_2$ labeled with Cy5, with the inset showing the corresponding mass (calc. 7361.8 Da; obs. 7360.9 Da, [M+TFA] 7472.2 Da, [M+2 TFA] 7584.1 Da).

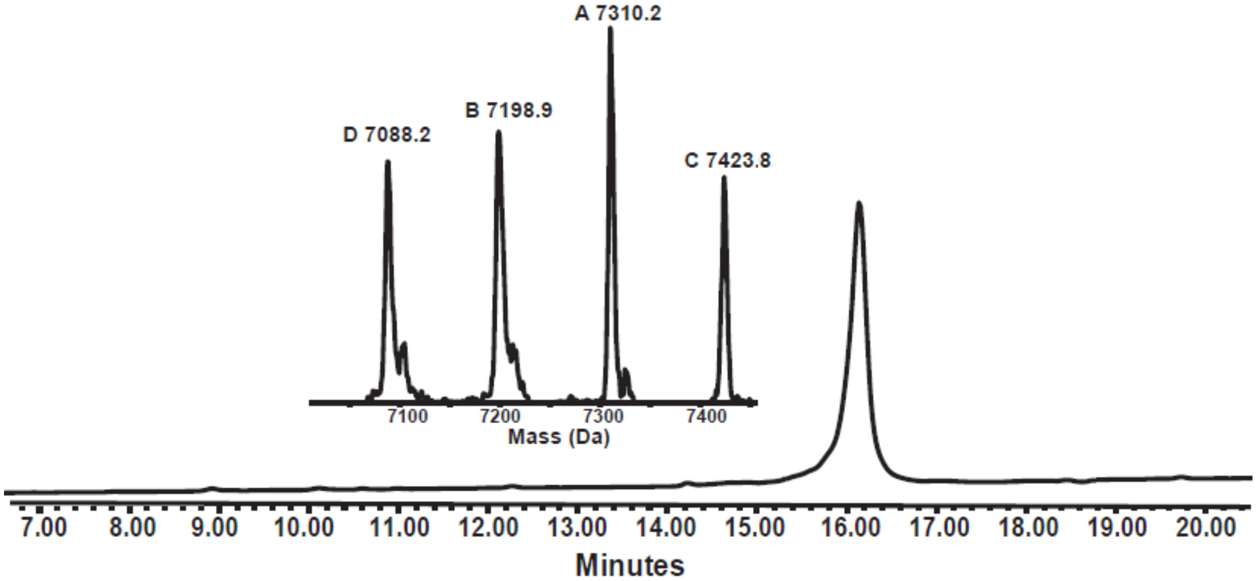


**Figure S15**. HPLC chromatograms and ESI-MS for purified *L*-Primordial-$(HhH)_2$-5G labeled with Cy5, with the inset showing the corresponding mass. (calc. 7089.3 Da; obs. 7088.2 Da, [M+1 TFA] 7198.9 Da] [M+2 TFA] 7310.2 Da, [M+3 TFA] 7423.8 Da).

**High-Resolution Mass Spectrometry of Proteins and Peptides Synthesized in this Work**

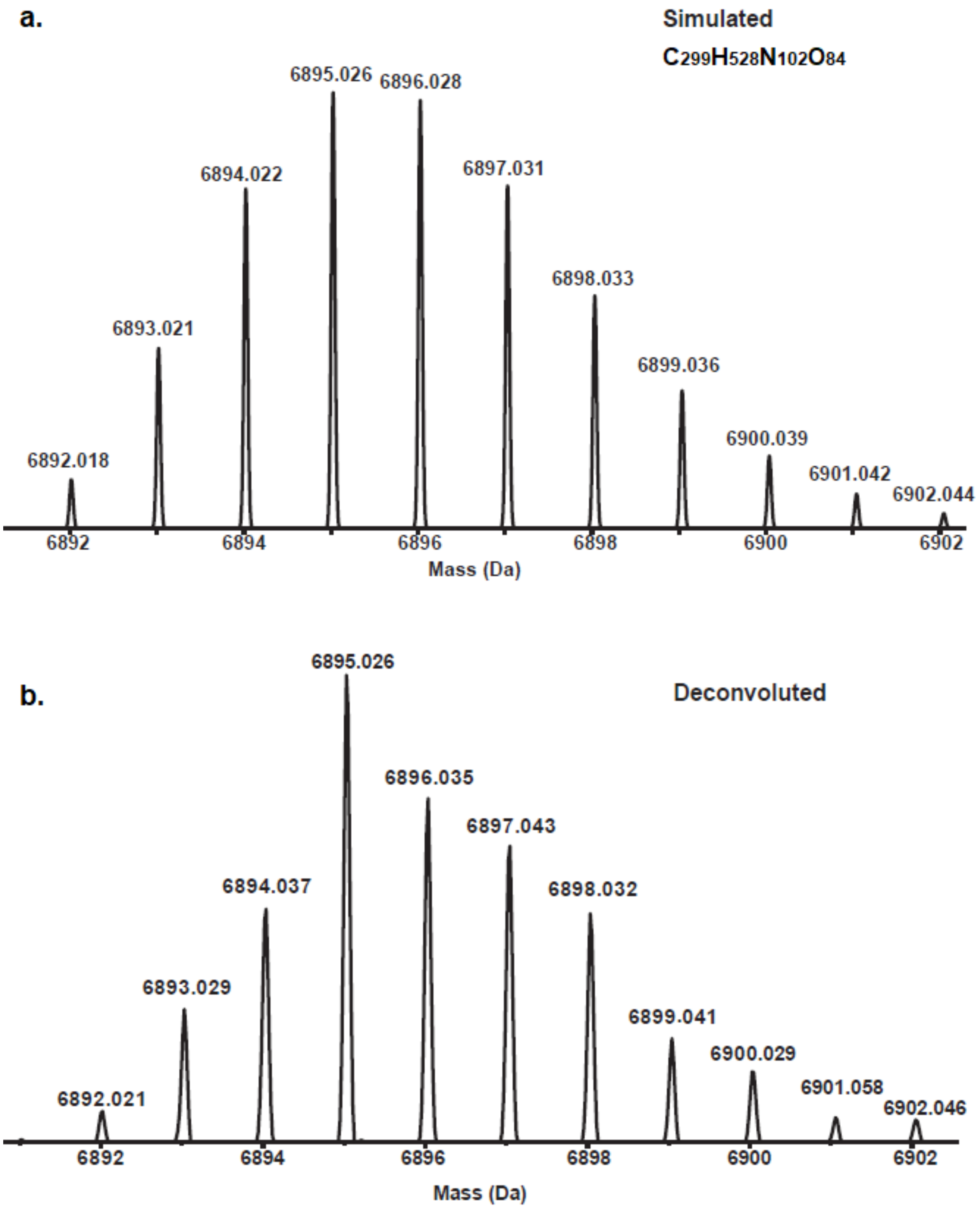


**Figure S16**. HR-MS analysis of *L*-Primordial-(HhH)$_2$. **a**. The simulated spectrum with chemical formula $C_{299}H_{528}N_{102}O_{84}$ is shown; **b**. The deconvoluted spectrum.

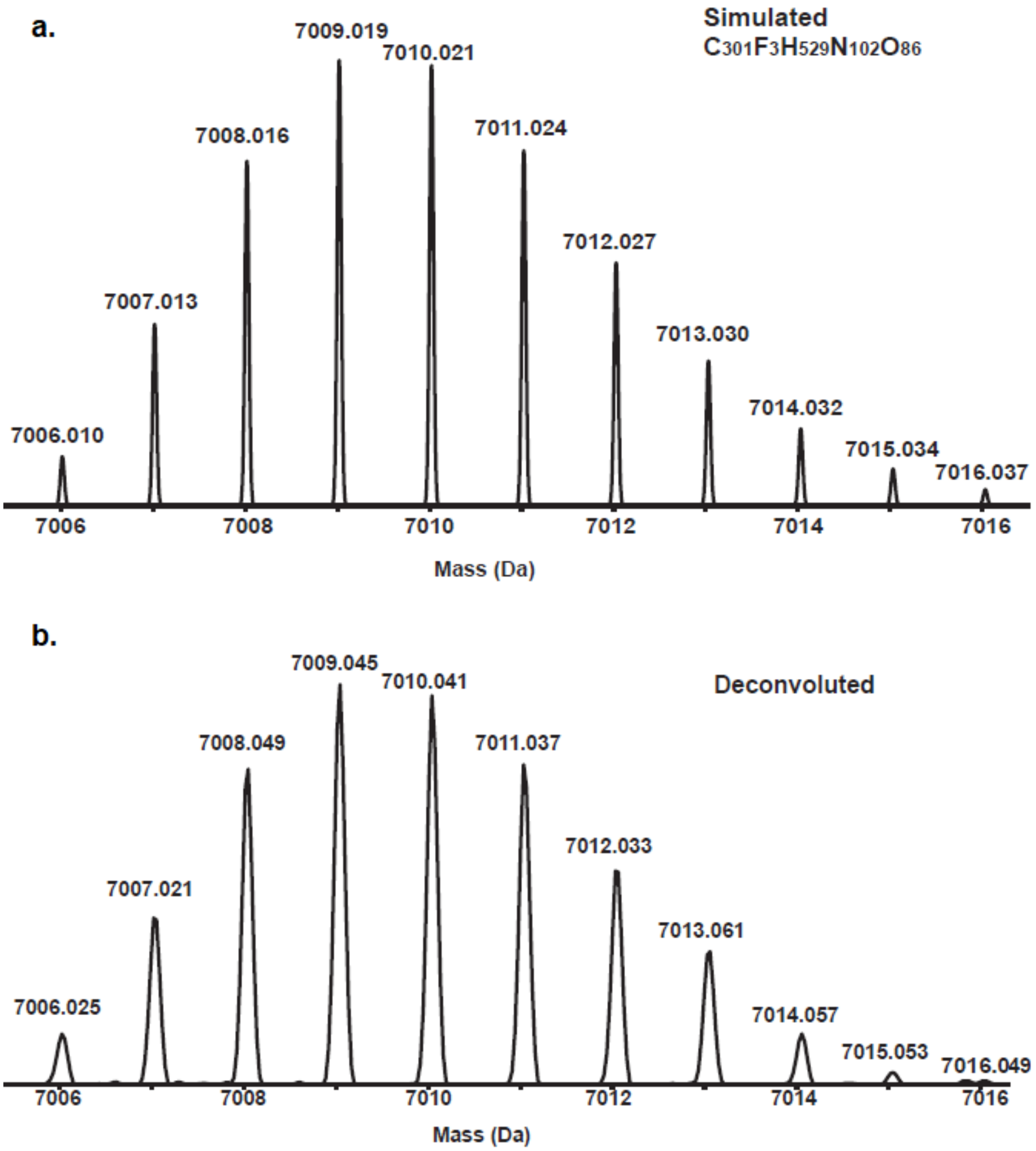


**Figure S17**. HR-MS analysis of *D*-Primordial-(HhH)$_2$. **a**. The simulated spectrum with one TFA molecule adduct, chemical formula $C_{301}F_3H_{529}N_{102}O_{86}$ is shown; **b**. The deconvoluted spectrum.

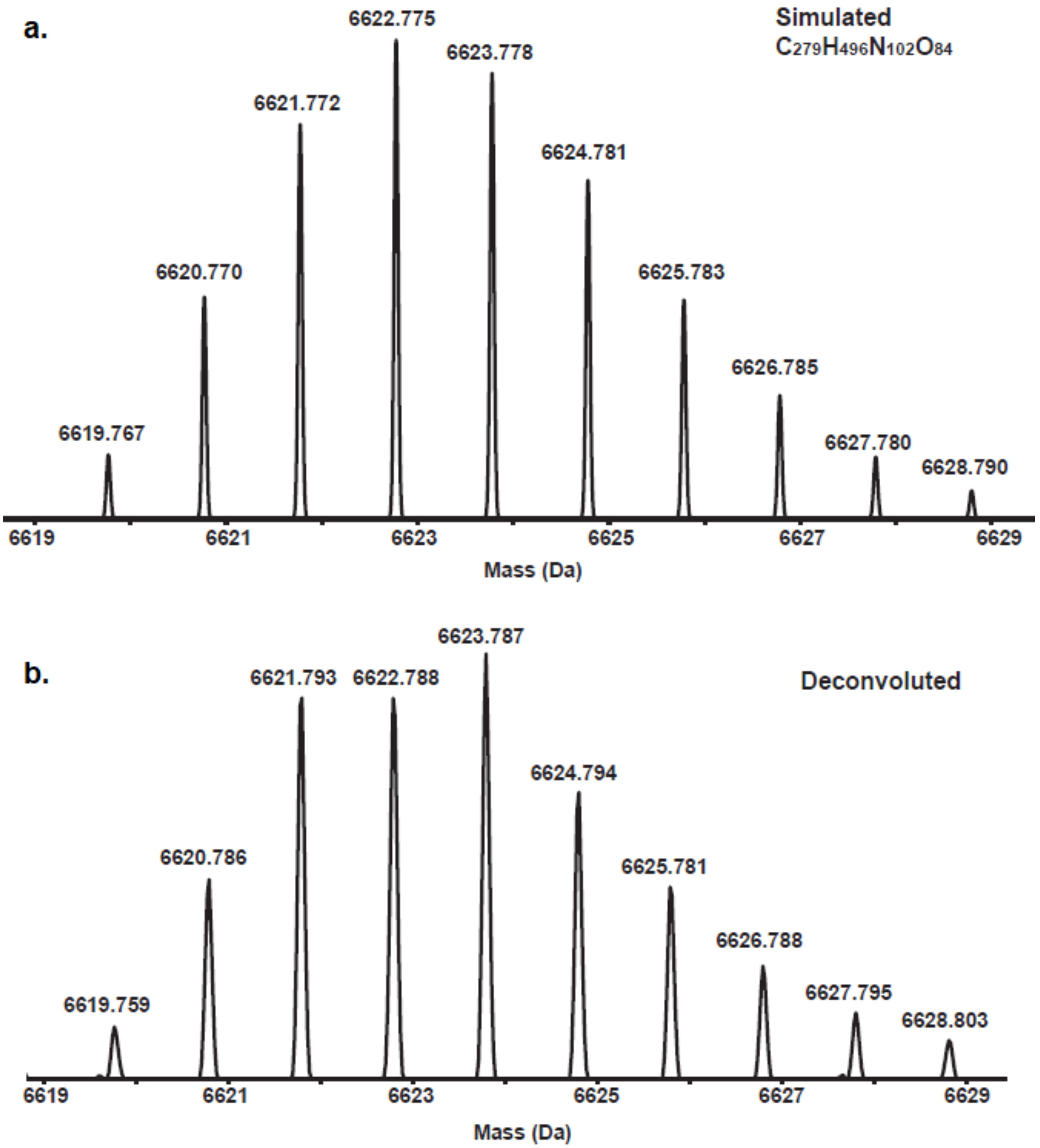


**Figure S18**. HR-MS analysis of *L*-Primordial-$(HhH)_2$-5G. **a**. The simulated spectrum with chemical formula $C_{279}H_{496}N_{102}O_{84}$ is shown; **b**. The deconvoluted spectrum.

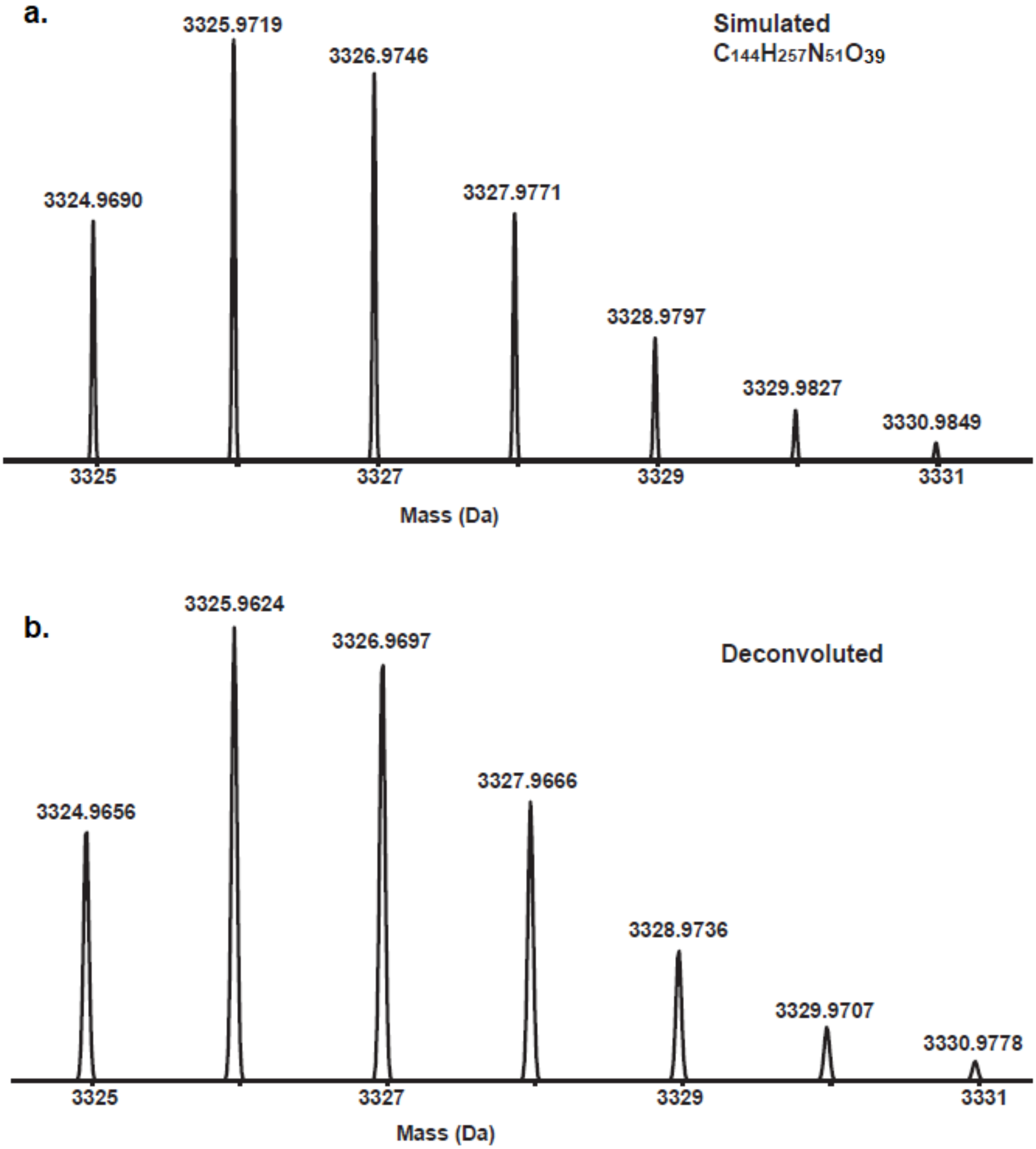


**Figure S19**. HR-MS analysis of *D/L*-Precursor-HhH. **a**. The simulated spectrum with chemical formula $C_{144}H_{257}N_{51}O_{39}$ is shown; **b**. The deconvoluted spectrum.

### Construct Characterization

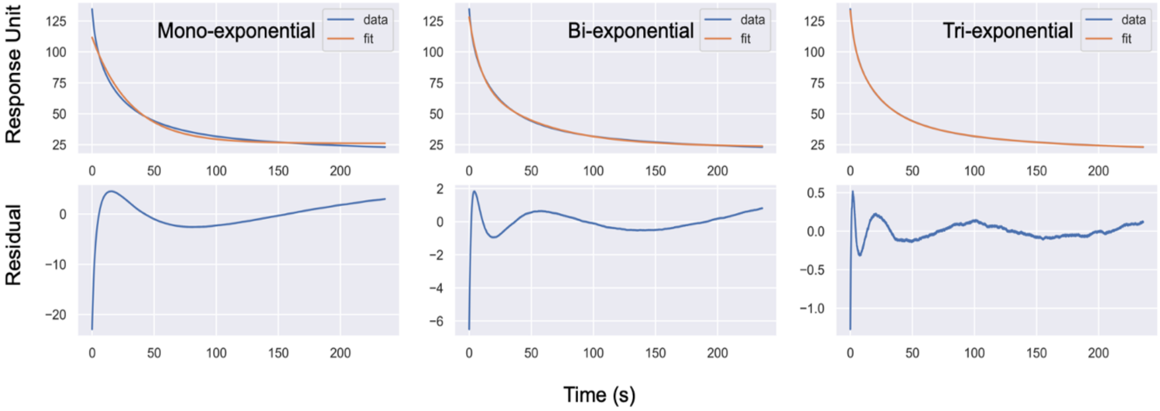


**Figure S20.** Representative dissociation curves. Within the window of useful SPR data, three kinetic phases can be reliably fitted. Although additional minor phases may be present, they could not be reliably fitted.

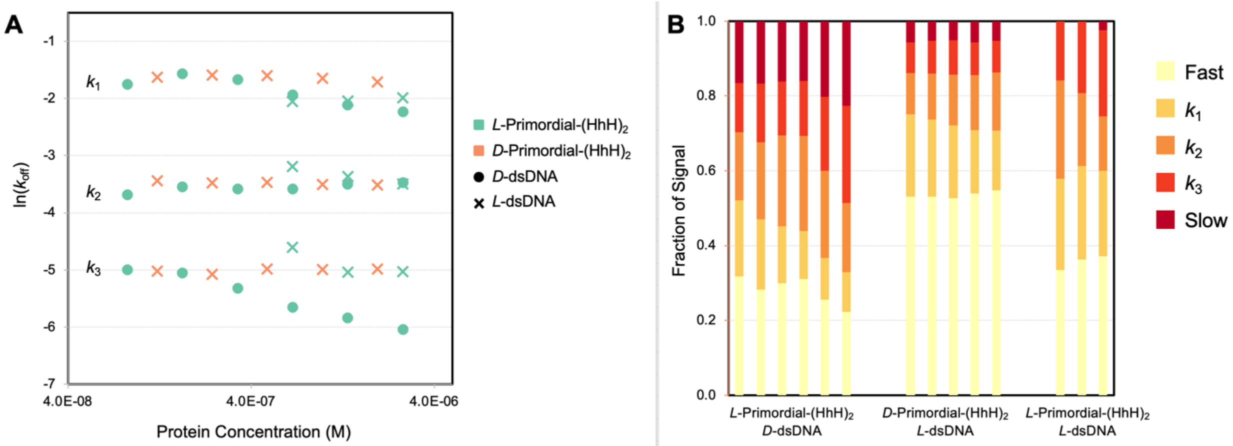


**Figure S21**. Analysis of SPR dissociation kinetics. **A** Kinetic phases associated with dissociation from dsDNA. As expected, off rates are largely concentration independent. Note that the dissociation rate constants associated with *L*-Primordial-(HhH)$_2$ binding to *D*-dsDNA is similar to the mirror image pair (*D*-Primordial-(HhH)$_2$ dissociating from *L*-dsDNA) and, surprisingly, also similar to the dissociation of *L*-Primordial-(HhH)$_2$ from *L*-dsDNA. **B**. The fraction of signal associated with each kinetic phase. Note that dissociation kinetics of the natural chiral pair are complex, and are associated with at least 5 kinetic phases: A fast phase (or phases) that occurs during a period of mechanical noise (due to needle movements) and high error (due to a refractive index change of the buffer); three kinetic phases that can be accurately modeled (see **Figure S20**); and a slow phase (or phases) that manifest as a positive shift in the baseline (and residual signal at t = ∞) but that can be eluted with 2M NaCl. The three phases that can be accurately modeled are relatively concentration independent. However, at higher concentrations of *L*-Primordial-(HhH)$_2$, the slowest of these three kinetic processes becomes slightly slower. This change may be due to cooperative binding effects as the dsDNA becomes progressively more coated with protein molecules that can then start interacting with each other and/or stabilize the optimal conformation of the dsDNA for binding. The flux through each kinetic mode associated with the mirror image pair are high similar to the natural pair, as expected, though with a notably higher contribution of the fast phase and a lower contribution of the slow phase. This difference is likely due to the lower synthetic purity of the *D*-protein and the *L*-RNA. Remarkably, the three measurable kinetic phases associated with *L*-protein binding to *D*-dsDNA are retained in *L*-protein binding to *L*-dsDNA. This conservation of kinetic phases may suggest a conservation of some binding modes. However, in the case of *L*-protein binding to *L*-dsDNA, the slow kinetic phase was almost completely abolished, perhaps suggesting the loss of the most stable binding mode.

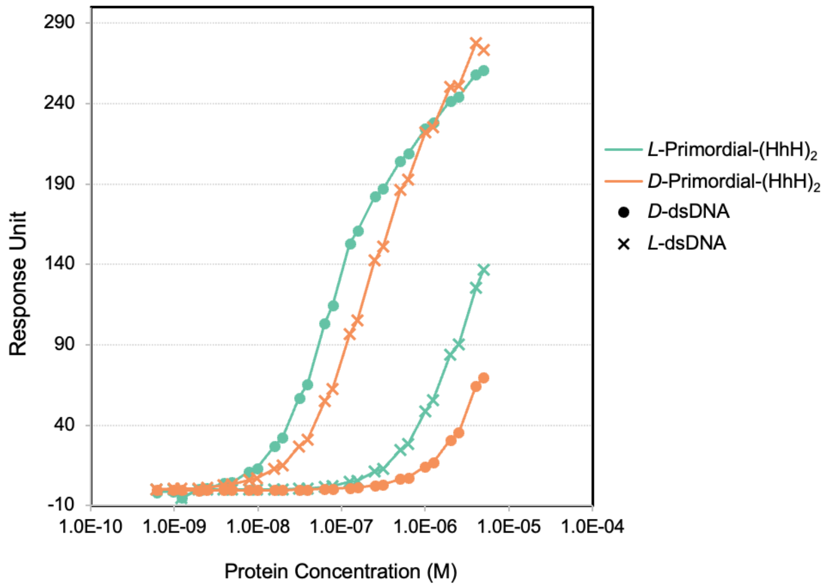


**Figure S22**. SPR stead state analysis of *L*- and *D*-Primordial-$(HhH)_2$ binding to *D*- and *L*-dsDNA. Note that these data were collected on a different SPR chip than that presented in **Figure 3** of the **Main Text**. As in **Figure 3**, stead state binding was estimated after 216 seconds injection.

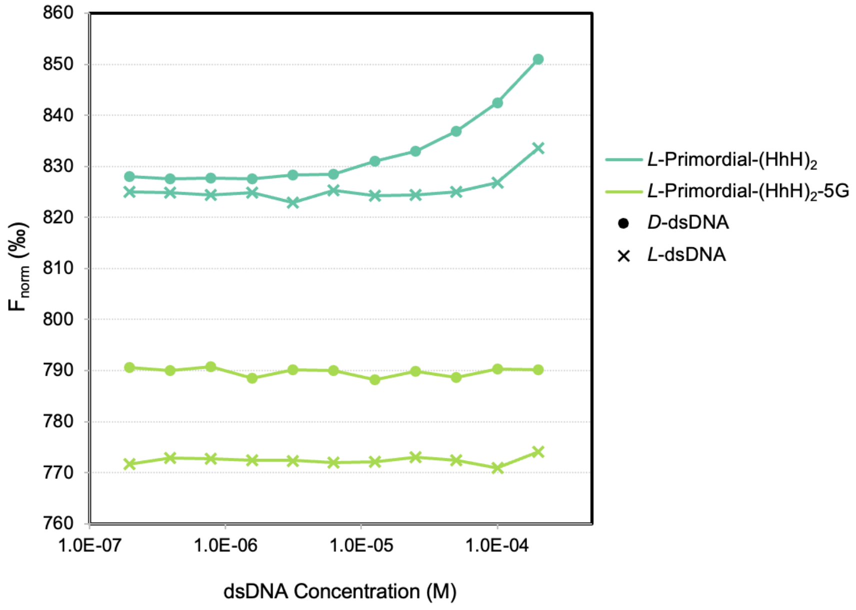


**Figure S23**. Microscale thermophoresis analysis. *L*-Primordial-(HhH)$_2$-5G, which has a disrupted PGIGP motif, does not bind to dsDNA of either chirality. *L*-Primordial-(HhH)$_2$, on the other hand, has indications of binding for both *L*- and *D*-dsDNA. Unfortunately, higher concentrations of ligands could not be analyzed due to significant changes in fluorescence of the conjugated Cy5 dye.

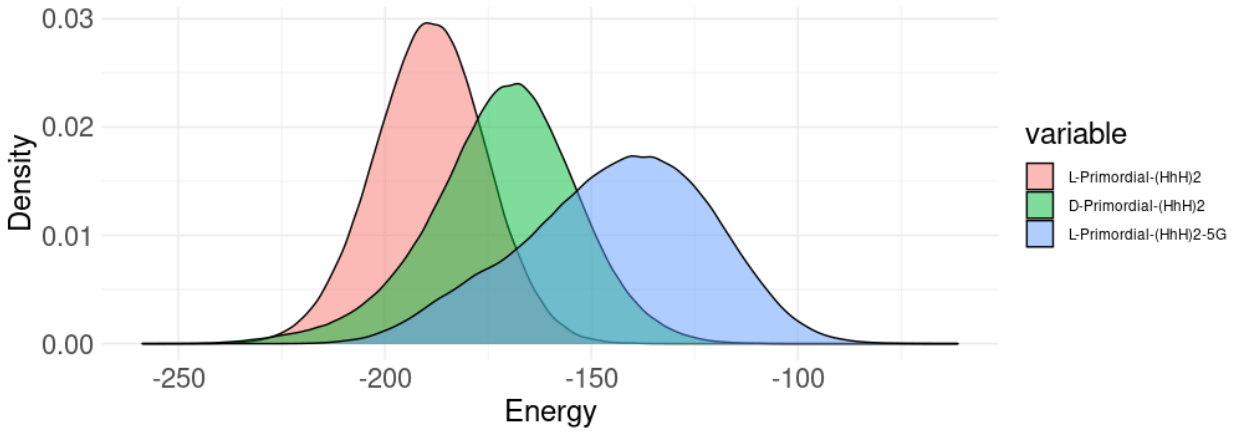


**Figure S24**. Stability distributions of Primordial-$(HhH)_2$ complexes with *D*-dsDNA after 500 ns of equilibration. LJ-SR energies calculated by the GROMACS energy method. Results are the mean of three independent simulations.

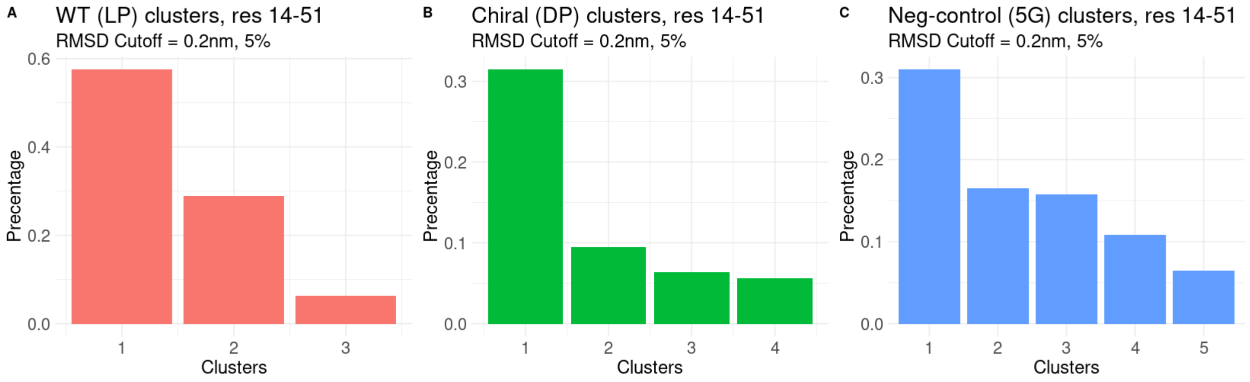


**Figure S25**. Clustering of protein-dsDNA complexes. Clustering of protein-dsDNA complexes. Clusters of similar conformations were identified using an RMSD cutoff of 0.2 nm for amino acids 14-51, excluding terminal tails. Only clusters representing at least 5% of total frames were considered.

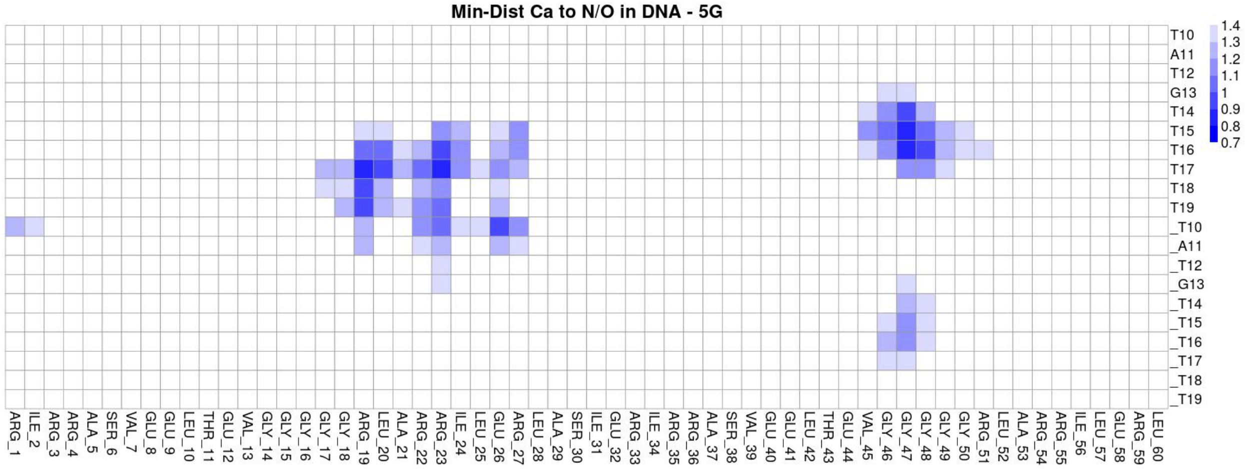


**Figure S26**.Heatmaps showing the minimum Cα to nitrogen or oxygen distances between protein and DNA, respective, for the dominant binding cluster of *L*-Primordial-(HhH)$_2$-5G. *Cf.* **Figure 4A**. The binding mode of *L*-Primordial-(HhH)$_2$-5G is significantly disported relative to *L*-Primordial-(HhH)$_2$.

**Table S1**. DNA sequences.

| **Experiment** | **Sequence** |
|---|---|
| SPR and MST<br>*D*- and *L*-dsDNA<br>(**Figure 3C-E**, **S22, S23**) | 5’biotin-CCGTCCGTAATCATGGTCATAGCTGTTTC-3'<br>(Reverse complement strand not biotinylated) |
| MD<br>*D*-dsDNA<br>(**Figure 4, S24**) | 5’-CGCTAGATCGATCGCTAGATC-3’ |
